# Design of a Synthetic sRNA-based Feedback Filter Module

**DOI:** 10.1101/504449

**Authors:** Nicolas Delalez, Aivar Sootla, George H. Wadhams, Antonis Papachristodoulou

## Abstract

Filters are widely used in engineering to reduce noise and/or the magnitude of a signal of interest. Feedback filters, or adaptive filters, are preferred if the signal noise distribution is unknown. One of the main challenges in Synthetic Biology remains the design of reliable constructs but these often fail to work as intended due, e.g. to their inherent stochasticity and burden on the host. Here we design, implement and test experimentally a biological feedback filter module based on small non-coding RNAs (sRNAs) and self-cleaving ribozymes. Mathematical modelling demonstrates that it attenuates noise for a large range of parameters due to negative feedback introduced by the use of ribozymes and sRNA. Our module modifies the steady-state response of the filtered signal, and hence can be used for tuning the feedback strength while also reducing noise. We demonstrated these properties theoretically on the TetR autorepressor, enhanced with our sRNA module.

## 1 Introduction

Synthetic Biology aims to design new or re-design existing biological devices and systems for a particular purpose. Examples include the design of ‘cellular factories’ producing valuable chemical compounds, biosensors capable of detecting toxins or viruses in a cell culture [Brophy and Voigt, 2014, Purnick and Weiss, 2009, Freemont and Kitney, 2015], or drug delivery systems [Zhou, 2016, Ozdemir et al., 2018]. Exploiting the intracellular machinery allows the synthesis of organic compounds that cannot be easily produced by other means, leading to novel applications in biotechnology, bioprocess engineering and cell-based medicine. However, one of the main challenges in Synthetic Biology remains the design of genetic systems that can be implemented in a predictable and robust way. Due to uncertainty, noise, burden and cross-talk inherent to biological systems, synthetic circuits can fail to work as intended. Indeed, elevated levels of protein production induce a high burden on the cell, notably by sequestering resources for transcription and translation (e.g. RNA polymerases and ribosomes) [Ceroni et al., 2015]. Operating at elevated protein production levels can also increase variability in the protein production due to intrinsic noise. To avoid these issues, common strategies to reduce the level of protein expression are to reduce the strength of promoters, the efficiency of the ribosome-binding site (RBS) or the plasmid copy number. However, transcriptional control is generally system dependent, diminishing the reliability of these approaches.

Filtering techniques are often used in signal processing, feedback control theory and communication systems to reduce signal noise [Haykin, 2002]. Filters can be classified into feedforward and feedback (or adaptive) filters. Feedforward filters are generally used when the noise statistics are known or can be estimated *a priori*; their output is the difference between the signal of interest and a modification of the same signal. Feedback filters automatically adjust their behaviour by comparing the output signal to the signal of interest at the input of the filter and thus are more favourable for signals corrupted by unknown noise distributions. In the context of Systems and Synthetic Biology, filtering capabilities of signalling cascades [Hooshangi et al., 2005, Thattai and van Oudenaarden, 2002], annihilation motifs [Laurenti et al., 2018] and other motifs [Samoilov et al., 2002] were studied in *silico*. Feedforward band-pass filters, which pass the signal only in a specific band of frequencies, have been constructed in *vivo* [Sohka et al., 2009], [Muranaka and Yokobayashi, 2010], while a noise attenuating feedforward filter was proposed and implemented in *vitro* in [Zechner et al., 2016].

The design of feedback filters is often performed with the help of feedback control theory, which has proven useful to render uncertain systems more reliable and robust to perturbations [Åström and Murray, 2008, Del Vecchio and Murray, 2015, Iglesias and Ingalls, 2010]. In a feedback loop, the output signal is measured and then used to modify the input of the system. In the filter design case, the controlled system is trivial: the signal corrupted by noise. Feedback control theory methods have been successfully applied in synthetic biology previously [Steel et al., 2017b], [Hsiao et al., 2018], [Ang and McMillen, 2013], [Briat et al., 2016], [El-Samad et al., 2002], [Del Vecchio et al., 2008], [Lillacci et al., 2018], [Cantone et al., 2009].

For example, in order to achieve a desired protein expression level an external computer was used to decide the input to the system (chemical or light induction) based on output measurements [Menolascina et al., 2011, Milias-Argeitis et al., 2011, Uhlendorf et al., 2012]. Such systems have inherent drawbacks, as control is achieved by interfacing the living cells with a digital computer that implements the control system.

Over the past few years, focus has shifted towards designing self-contained *in vivo* controllers. While the vast majority of these experimental implementations were protein-based [Hsiao et al., 2014, Folliard et al., 2017, Rosenfeld et al., 2002] small non-coding RNAs (sRNAs) have also been recently used in this context [Ghodasara and Voigt, 2017, Takahashi et al., 2014, Hu et al., 2018, Kelly et al., 2018]. sRNAs are found in all domains of life and have been shown to play critical regulatory roles in many processes [Cech and Steitz, 2014], [Michaux et al., 2014], [Robledo et al., 2018], [Gottesman and Storz, 2011], [Livny and Waldor, 2007], [Nitzan et al., 2017]. Most sRNAs characterised to date act as post-transcriptional regulators by interacting with specific mRNA targets. Feedback loops involving sRNAs can be found in natural biological processes, for example in the regulation of the expression of quorum-sensing genes [Liu et al., 2013] and in the promotion of a switch for adequate Lrp-dependent adaptation to nutrient availability [Holmqvist et al., 2012]. Post-transcriptional down-regulation is favourable since no proteins are being expressed in this regulation mechanism. Instead, sRNAs are produced quickly, potentially propagating signals rapidly [Holmqvist et al., 2012, Hussein and Lim, 2012, Mehta et al., 2008, Takahashi et al., 2014] and require less energy than proteins, hence reducing the burden to the host. Their operational dynamics are also much faster due to their naturally high degradation rate [Hussein and Lim, 2012]. Therefore, sRNAs provide a promising alternative to the commonly used transcriptional control [Steel et al., 2017, Agrawal et al., 2018].

In this work, we considered two sRNA-based designs to filter variations in transcription, shown in Figure 1. In the first design the regulatory sRNA is placed under the control of a *separate* promoter to the one controlling transcription of a target gene (henceforth *in trans* design). In the second design, the sRNA is placed directly downstream of the target gene *in cis* so that both are under the control of the *same* inducible promoter. The *in cis* design also contains a self-cleaving ribozyme between the regulated mRNA and the sRNA sequences, as experiments demonstrated that the mRNA-sRNA strand needs to be separated for the translational attenuation to be efficient. We computationally showed the benefits of the *in cis* in comparison to the *in trans* design. While modelling the two circuits and performing numerical simulations showed that the mean steady-state values in both design are attenuated at similar levels, it was evident that the *in cis* design reduces noise significantly, while the *in trans* design can adversely amplify it. Modelling also revealed that the *in trans* design operates approximately as a feedforward filter, in that its output is the mRNA available after sRNA regulation while the *in cis* design also contains a feedback component, in that the free sRNA produced by self-cleavage of the ribozyme can regulate the amount of mRNA-sRNA transcript available for cleavage.

**Figure 1:**
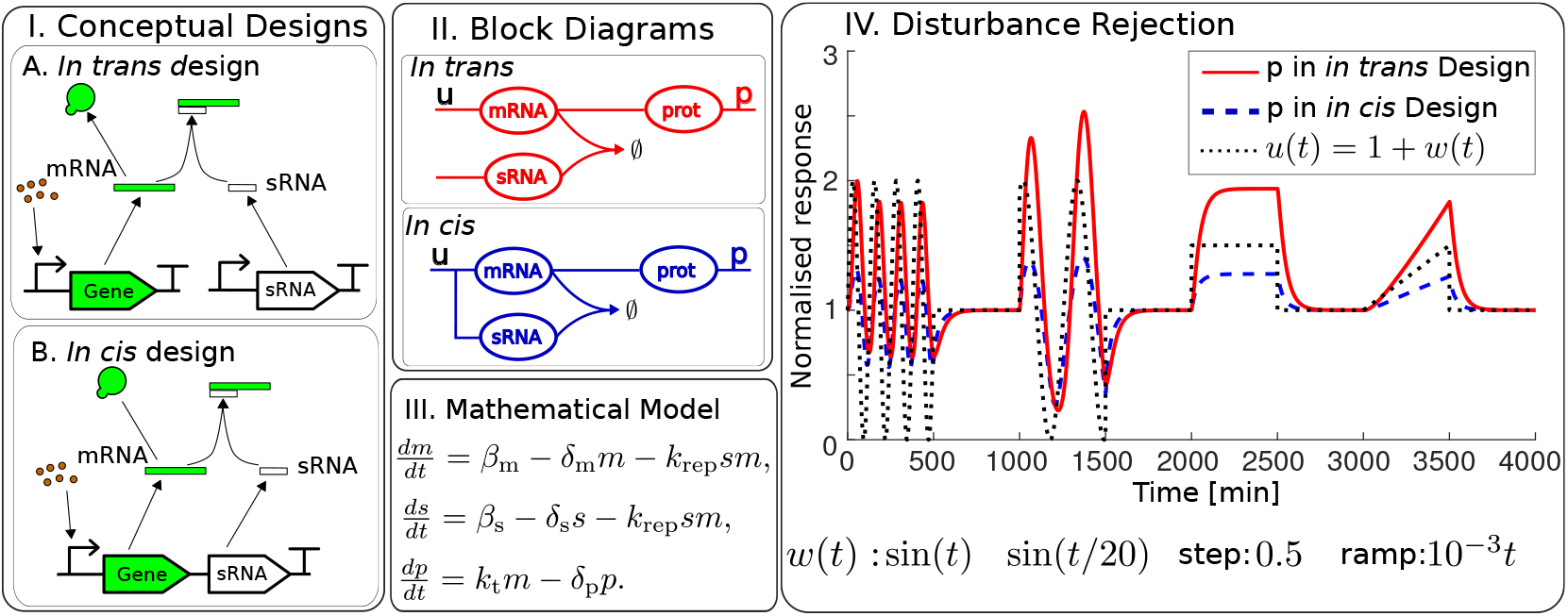
I. Two conceptual designs of filters using sRNA. In the in *trans* design, the mRNA and the sRNA are under two separate promoters, while in the *in cis* design the sRNA is placed downstream of the mRNA under the same promoter. II. Block diagrams of the *in trans* and *in cis* designs. III. Mathematical model for the designs. In the *in cis* design, we have *β_m_* = *β_s_* = *β_ms_*, where *β_ms_* is the production rate of mRNA and sRNA in the in *cis* design. IV. Improved disturbance rejection in the *in cis* design. In both designs, the perturbation was applied on the mRNA production rate. In the in *trans* design, *β_m_* = u(t) = 1 + w(t) [nM/min] and in the *in cis* design *β_ms_* = 1 + w(t) [nM/min]. In the plot, the protein concentrations were normalised by dividing by the steady-state expression of the models with *β_m_* =, *β*_ms_ = 1 [nM/min]. The simulations show that the signal w(t) is attenuated more efficiently in the in cis design than the *in trans* design. These simulations also indicate that the transcription noise should be attenuated more efficiently in the *in cis* design.

As the *in cis* design also attenuates the mean steady-state of the signal, this module can also be used in feedback control in order to reduce the strength of the feedback. We demonstrate the value of the *in cis* design on the P_tet_/TetR autorepressor. Here, sRNA is used to tune the TetR feedback strength without modifying the rest of the system. Our numerical simulations suggest that the *in cis* design offers a tunable response in terms of the mean output while attenuating transcription noise.

## 2 Results

### 2.1 Conceptual designs of sRNA-based filters

We first considered the conceptual designs of the *in trans* and *in cis* filters depicted in Figure 1.I (and as block diagrams in Figure 1.II), which can be modelled using a similar set of reactions. In the *in trans* design we assumed the following reactions: 
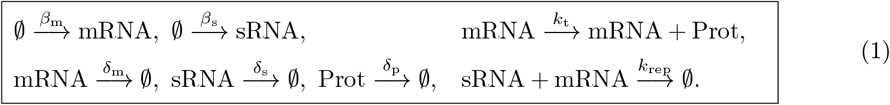
where Prot denotes a protein, which is the filter output. In this design, mRNA and sRNA are transcribed in two different chemical reactions with rates *β_m_*, *β_s_*, respectively.

In the *in cis* design, however, mRNA and sRNA are transcribed in the same reaction with the same transcription rate *β_ms_*, so that this model takes the form 
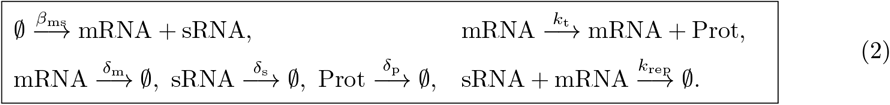

This assumes that the transcribed strand containing mRNA and sRNA splits into mRNA and sRNA instantaneously. This conceptual (or ideal) representation drove the biological implementation discussed later in the text.

In both designs we assumed that mRNA is translated into a protein at the rate *k_t_* and that the degradation/dilution rate for every species is different, as mRNA generally degrades faster than proteins and the reported values for the degradation rate of sRNA vary [Hussein and Lim, 2012]. We also assumed that the rate of mRNA-sRNA unbinding is negligibly small, as previously reported [Hussein and Lim, 2012, Kelly et al., 2018], and therefore we did not include it in our model. Modelling both designs using mass-action kinetics yielded the model presented in Figure 1.III, with the difference that for the *in cis* design, we have *β_m_* = *β_s_ = β_ms_*.

Note that while sRNA down-regulates the translation process in both designs, the two designs lead to different responses to disturbances in the mRNA transcription process. Indeed, the *in cis* design should be able to attenuate the transcription disturbance better since for every molecule of mRNA produced, so is one molecule of sRNA. Therefore, a burst in transcription of mRNA would also result in a burst in transcription of sRNA. To illustrate the response to disturbances in transcription, we varied the production rate of mRNA in both systems simultaneously, that is we used *β_m_* = *β_ms_* = *u*(*t*) = 1 + *w*(*t*)[nM/min], where *w*(*t*) is the disturbance signal (see Figure 1.IV for the used signals *w*(*t*)), and we set *β_s_* = 1 [nM/min], *k*_rep_ = 0.5 [1/(nM min)],*δ*_m_ = 0.2476 [1/min], *δ*_S_ = 0.0482 [1/min], *δ*_p_ = 0.0234 [1/min] and *k_t_* = 1 [1/min] (see Table S1 in SI). The results of the simulation are shown in Figure 1.IV, where the protein concentrations with *β_m_* =, *γ*_ms_ = *u*(*t*) were divided by the steady-state protein concentrations with *β_m_* =, *γ_ms_* = 1 giving the normalised response. The results clearly indicate that the in cis design attenuates the disturbance better than the *in trans* design.

### 2.2 Biological implementation of the *in cis* design

#### 2.2.1 Importance of mRNA-sRNA cleavage in the *in cis* design

Next we constructed the *in cis* design in the laboratory and to test experimentally whether controlled attenuation could be achieved using this design. To further minimize the burden on the cell, we chose to use a low copy number plasmid as the vector to implement our *in cis* RNA-based attenuator design (Table S4 in SI). We also chose to use *P_tet_* as the inducible promoter as it offers tight regulation in response to aTc. As a proof-of-principle, we chose sfGFP as the output to be attenuated. The synthetic regulatory sRNA was designed following the protocol described by [Na et al., 2013, Yoo et al., 2013], in which we changed the binding sequence to target *sfgfp*. The sequence of our construct hence consists of an *sfgfp*, ribozyme, the synthetic sRNA consisting of the target binding sequence (TBS) followed by an Hfq-recruiting *micC* scaffold. Based on [Yoo et al., 2013], we chose a 25-nucleotide long sequence as a starting point for the TBS. Using the web-based service DINAMelt, this sequence gave a Δ*G* = −30.4 kcal • mol^−1^, in line with full translation inhibition in [Yoo et al., 2013]. We also hypothesized that the sRNA should be cleaved off the mRNA strand for efficient binding and translation inhibition. We therefore introduced a self-cleaving ribozyme, the Human Hammerhead Ribozyme 9 (HHR9) shown to work well *in vivo* [De la Peña et al., 2003, De La Peña and García-Robles, 2010, Perreault et al., 2011], between *sfgfp* and the sRNA.

We monitored cell fluorescence over time in response to varying levels of aTc for two constructs, one with no ribozyme and one carrying HHR9, and compared them with the fluorescence from cells lacking the ribozyme/sRNA part.Figure 2.II shows the steady-state levels of normalized fluorescence for each strain. Attenuation of the output is only observed for the construct expressing the HHR9 ribozyme, confirming our hypothesis that cleavage of the sRNA from the target mRNA is necessary for efficient translation inhibition.

**Figure 2:**
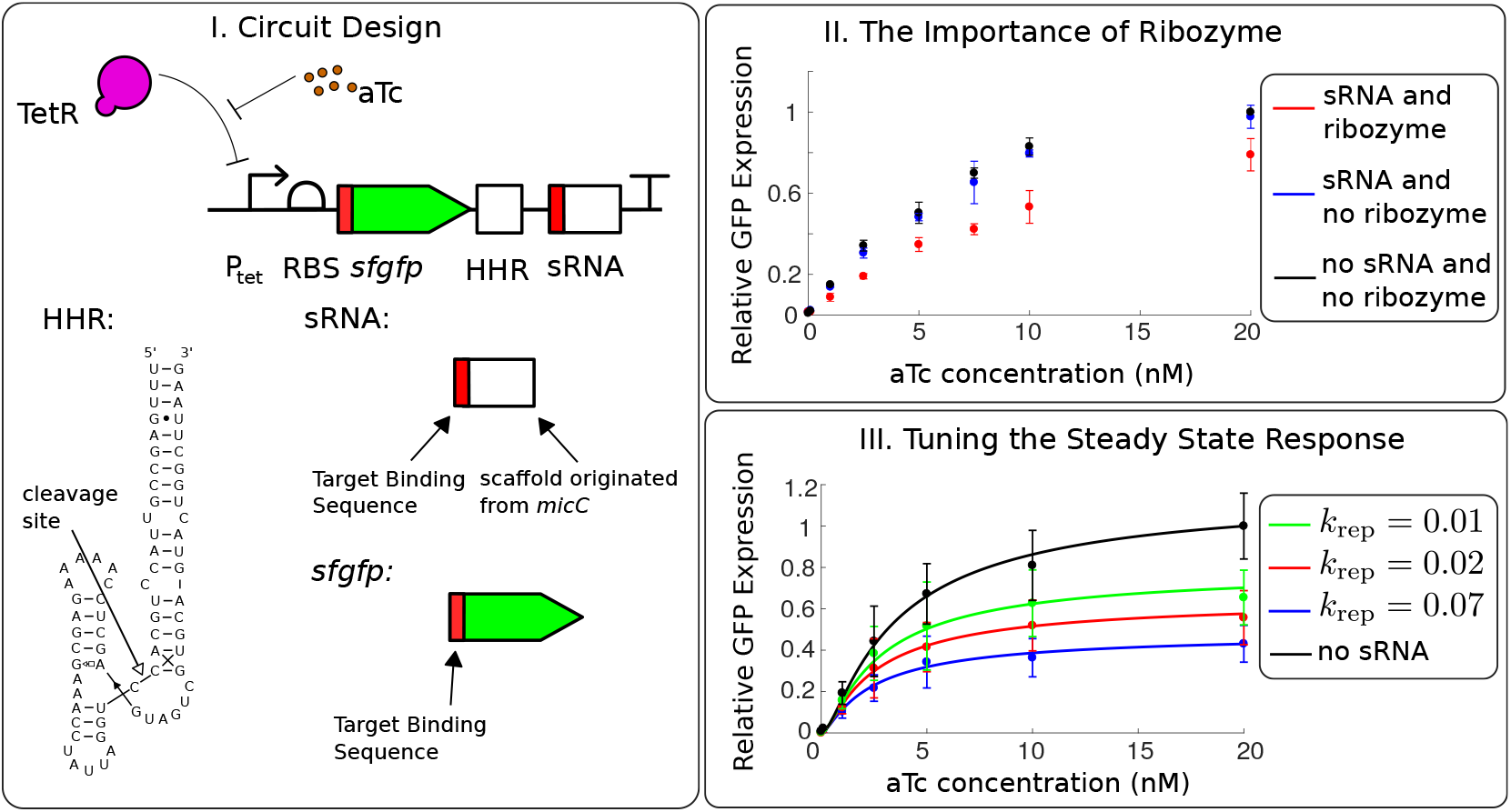
Implementation of the in *cis* filter. I. Experimental design: sfGFP and sRNA are placed under the control of a *P_tet_* promoter and separated by an HHR9 ribozyme. The schematics for the ribozyme and the synthetic sRNA are adapted from [Perreault et al., 2011] and [Yoo et al., 2013], respectively. II. Importance of the ribozyme for efficient attenuation. Fluorescence output measured at different aTc concentrations for designs with and without HHR9, compared to the fluorescence of a system without sRNA attenuation. III. Fine tuning the attenuation level. Fluorescence output measured for varying length of the target binding sequence (TBS) at different aTc concentrations. Solid lines correspond to model predictions.

#### 2.2.2 Fine tuning the steady-state level

We next tested the possibility of fine tuning the level of attenuation by modifying the TBS of the sRNA, following the protocol described in [Yoo et al., 2013]. To do so, we decided to either increase or decrease the length of the TBS in the construct with the HHR9 ribozyme, leading to an increase and decrease of the translation inhibition, respectively. We estimated the different binding energies using DINAMelt and chose four different new sequence lengths to test: 30–, 27– and 22–nucleotides long, giving binding energies Δ*G* = –38.2 kcal • mol^−1^, Δ*G* = –31.6 kcal *·*mol^−1^, Δ*G* = –28.6 kcal *·*mol^−1^, respectively. We monitored the cell fluorescence over time in response to varying levels of aTc for each construct.Figure 2.III shows the output of the system for the different binding energies, displayed as the normalised fluorescence plotted against different aTc concentrations. The output can be reduced to 40% of the signal (for the longest TBS tested) and its value can be varied by altering the length of the TBS, as predicted.

### 2.3 Modelling and analysis of the *in cis* filter

#### 2.3.1 Modelling the *in cis* filter

Having established that the conceptual designs can be implemented experimentally, we proceeded with a more detailed mathematical model to understand further their properties. For convenience we labelled the mRNA of GFP as mGFP. We assumed that self-cleavage of the ribozyme takes place after transcription of the full RNA, that is, mGFP-ribozyme-sRNA (labelled fmRNA) is cleaved into mGFP and sRNA with a rate *k*_rc_. We assumed that sRNA binds to mGFP preventing GFP translation. We also assumed that sRNA can bind to fmRNA, which can then self-cleave into sRNA and an mGFP-sRNA complex. Since experimental data suggests that the presence of a ribozyme is essential for sRNA and mRNA binding in the *in cis* design, we assumed that fmRNA (mGFP-ribozyme-sRNA strand) does not bind to the target mRNA (mGFP). We assumed that GFP can be translated both from mGFP and fmRNA. The other reactions were assumed to be the same as for the *in trans* design, leading to the following chemical reaction model: 
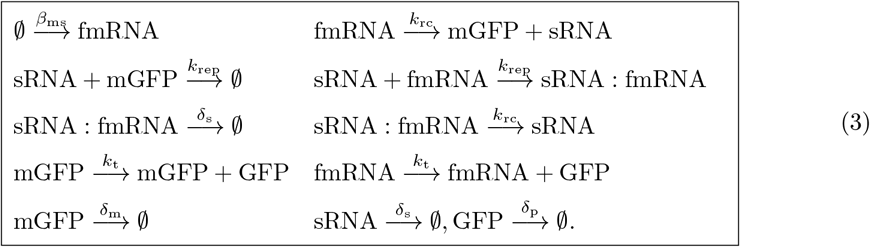

We followed the standard mass-action kinetics modelling framework and obtained the model presented in Figure 3.III. We analysed the resulting model as described in the SI. In particular, the frequency domain analysis showed that both *in cis* and *in trans* designs implement a low-pass filter attenuating high frequency noise. For realistic parameter values the term *k*_rc_ *c* — *k*_rep_ *sf* in Figure 3 remains close to zero, therefore, it does not significantly affect the equation of the sRNA concentration (s) and we hence pictorially represent that sRNA directly degrades fmRNA in Figure 3.I. Depicting the *in cis* design in the block diagram in Figure 3.II revealed the structure of the filter. The mGFP and sRNA interaction represents *the feedforward part* of the filter from the transcription initiation, since sRNA and mGFP are produced at similar time instances and sRNA binds to mRNA forming an inert complex. There is also *a feedback part* in this design formed by the sRNA and the fmRNA interaction. Indeed, fmRNA self-cleaves into mGFP and sRNA, which then binds to fmRNA forming the complex, which contains a ribozyme and splits to an inert complex mGFP-sRNA and a free sRNA.

**Figure 3:**
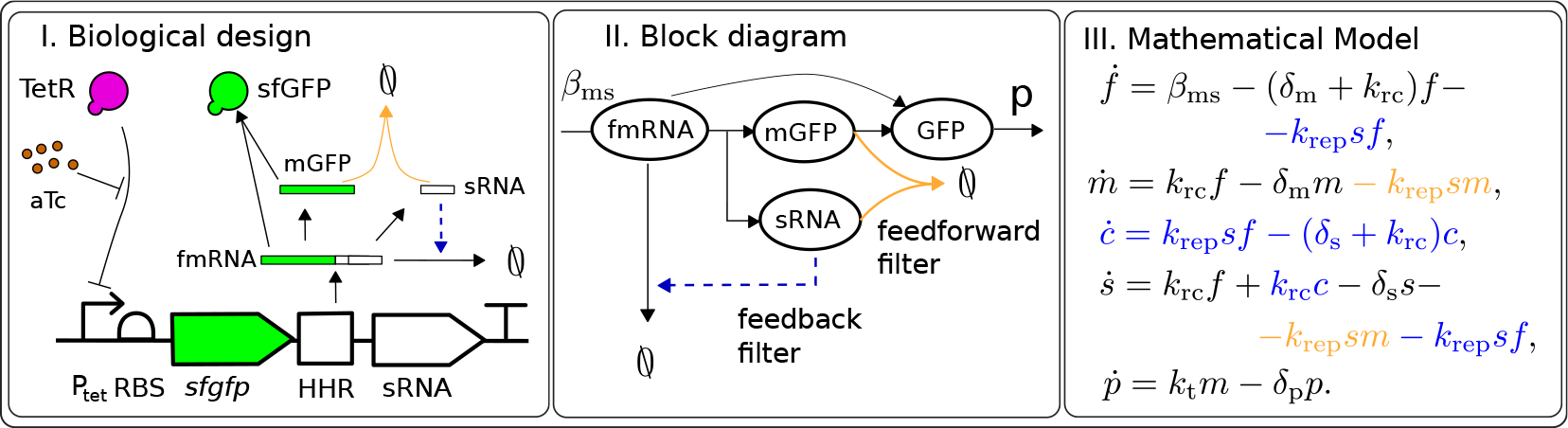
A biological implementation of the *in cis* design. fmRNA denotes the RNA strand containing mGFP (mRNA of GFP), ribozyme and sRNA. The sRNA-mGFP interaction forms the feedforward part of the filter, while the fmRNA-sRNA interaction forms the feedback part of the filter. The dashed line between sRNA to the fmRNA degradation signifies that sRNA binds to fmRNA forming a complex that cleaves into an inert mGFP-sRNA complex and another sRNA copy, thus the sRNA copy number does not decrease. II. Block diagram of the *in cis* design. III. A mathematical model of the *in cis* design, where *f*, *m*, *c*, *s*, and *p* denote the concentrations of fmRNA, mGFP, sRNA:fmRNA complex, sRNA and GFP, respectively. For the chosen parameter values, the term *k*_rc_ *c* — *k*_rep_ *sf* remains close to zero, therefore, it does not significantly affect the equation of the sRNA concentration (s) and we can assume that sRNA degrades fmRNA directly.

We also derived a non-dimensional model of the *in cis* design, which clearly exhibited timescale separation between the quantities *f* + *m*, *s, p* on one side and *f*, *c* on the other (see SI for details). This allowed the derivation of a simplified deterministic model of the *in cis* filter 
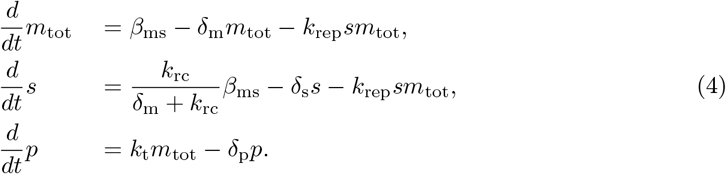
where *m*_tot_ is the total concentration of fmRNA and mGFP. The key assumption for this analysis was the faster ribozyme cleavage rate in comparison to other reactions. Further investigation revealed strong stability properties of the simplified model, in particular, we ruled out oscillations and multiple steady-states under some assumptions.

Our analysis suggests a possible tuning dial in the in cis design: the ribozyme (with cleavage rate k_rc_) can be used to adjust the gain of the output attenuation, as well as the sRNA-mGFP binding strength *k*_rep_. With the ribozyme cleavage rate increasing, the deterministic model for this system converges to the ‘ideal’ model of the conceptual design, however, the reported values of the ribozyme cleavage rate *k*_rc_ are not large enough for us to assume that 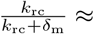 and so that we cannot discard the ribozyme cleavage rate completely. While the simplified model was useful for the analysis and revealed the mathematical difference between the in cis and in *trans* designs, it hid the feedback part of the filter. This raised the question if the feedback part of the filter has a significant effect on the repression of translation.

#### 2.3.2 *In silico* evidence of the feedback in the *in cis* design

Here we evaluated the influence of the feedback on the repression of translation. We performed model simulations of the in cis design, and the models of in cis design without the feedforward part (mGFP and sRNA binding) and without the feedback part (fmRNA and sRNA binding). We set *k*_rep_ = 0.5 [1/(nM min)], *k*_rc_ = 5 [1/min] *δ*_m_ = 0.2476 [1/min], *δ*_s_ = 0.048 [1/min], *δ*_p_ = 0.0234 [1/min] and *k_t_* = 1 [1/min] (see Table S1 in SI). We replaced the production rate of fmRNA *β*_ms_ in all three systems with 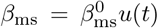, where *u*(*t*) is the disturbance signal depicted by dashed purple line in Figure 4.I and 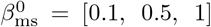 = [0.1, 0.5, 1] [nM/min]. We plot the response of the systems divided by the response with *u*(*t*) ≡ 1. Numerical simulations presented in Figure 4.I clearly suggest that the feedback part of the filter has a larger influence on the steady-state behaviour than the feedforward part even with a high ribozyme cleavage rate *k*_rc_ = 5 [1/min].

**Figure 4:**
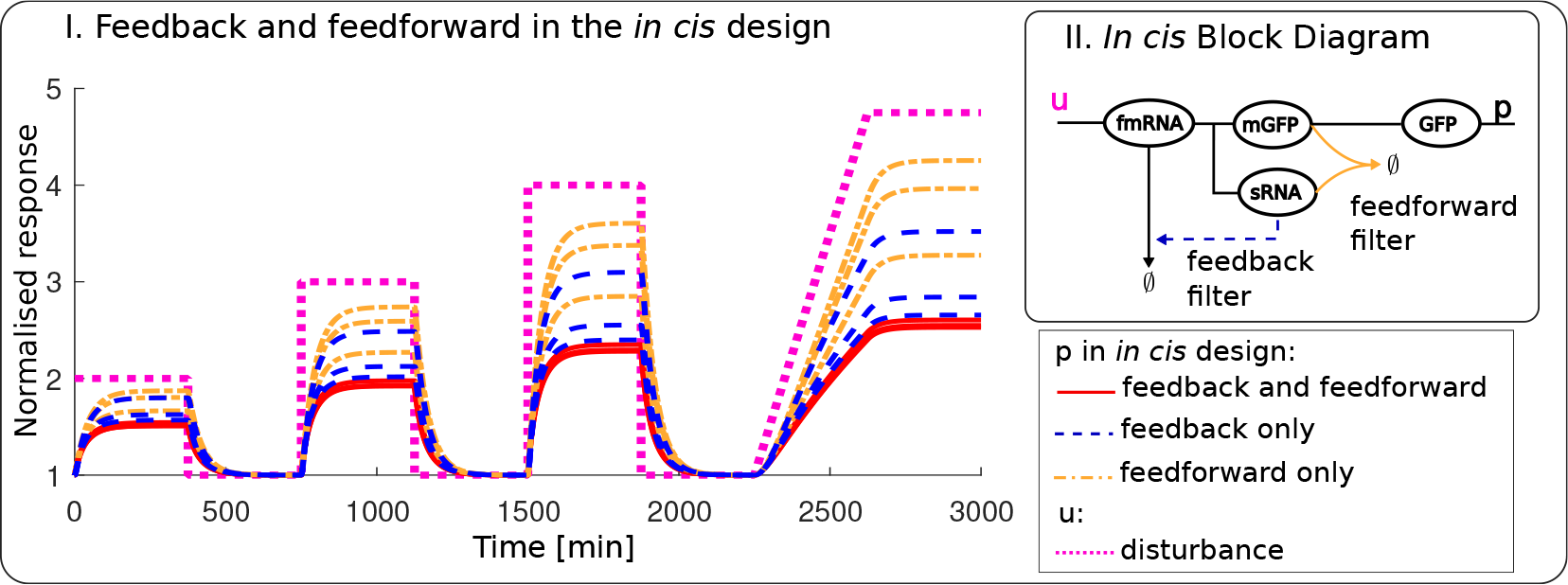
I. The significance of the feedback part of the filter validated by deterministic in *silico* analysis of the in *cis* design. The simulations are performed with and without feed-forward/feedback, and 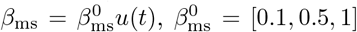 [nM/min]. Knocking out the sRNA feedback results in weaker signal repression than knocking out the sRNA feedforward. II. Block diagram of the in cis design.

### 2.3.3 *In cis* filter improves the noise properties of the signal

The simulations of the conceptual model suggest that the *in cis* design attenuates intrinsic noise of the promoter in a much more efficient way than the *in trans* design. We verified this hypothesis by performing stochastic simulations using the Gillespie Algorithm with the parameters/parameter ranges in Table S1. We considered the coefficient of variation as a noise metric (Figure 5). We plotted the coefficient of variation relative to the mean steady-state for each design. These numerical simulations suggest that the *in trans* design has a very narrow range of *k*_rep_ values for which noise is attenuated, when compared to the circuit with no sRNA (or *k*_rep_ = 0) while the *in cis* design attenuates noise for almost all values of k_rep_. An example is presented in the caption of Figure 5, while the numerical values are given in Table S2 in SI. This analysis suggests a simple method to design the *in cis* filter: choose the maximum possible combination of *β*_ms_, *k*_rep_ that achieves the desired GFP mean values.

**Figure 5:**
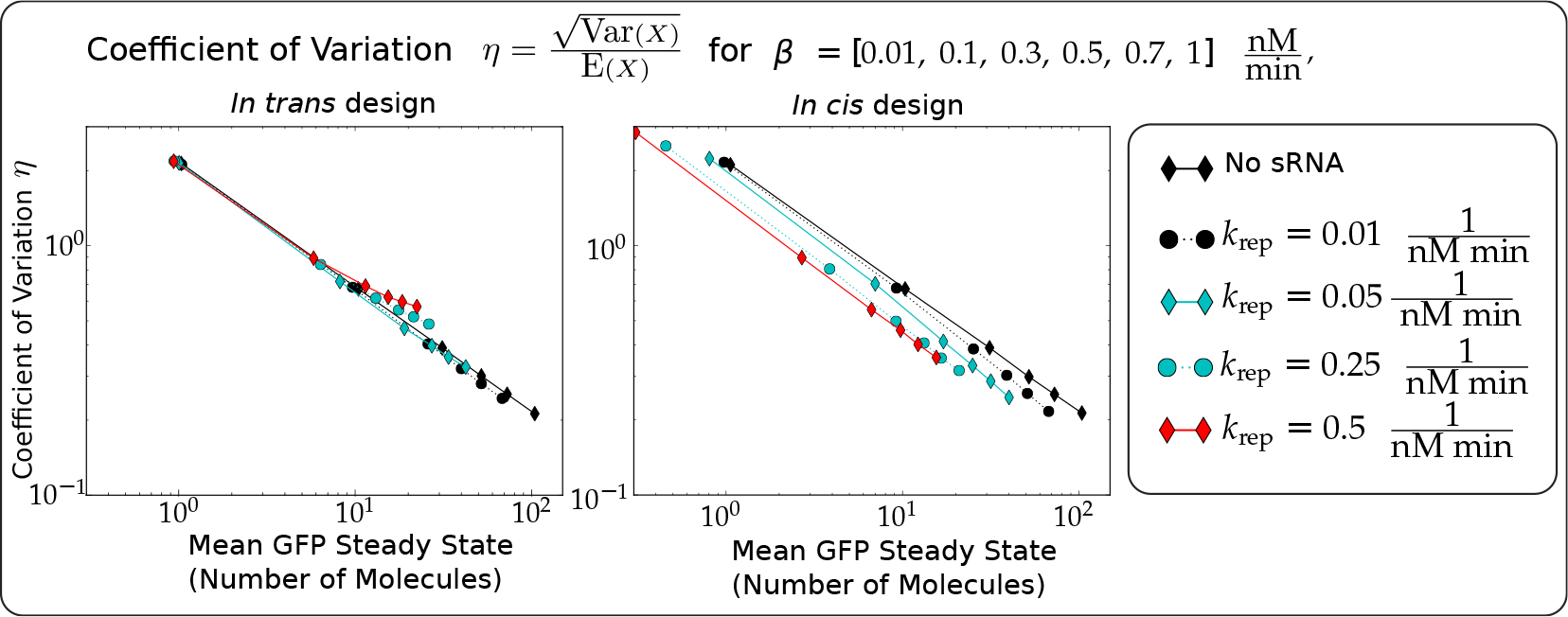
*In cis* filter effectively reduces noise in comparison to *in trans* filter. Data points for each line correspond to different values of *β* from [0.01, 0.1, 0.3, 0.5, 0.7, 1] [nM/min]. Every line corresponds to a different repression sRNA-mRNA binding strength *k*_rep_. For the *in trans design* we set *β*_m_ = *β*_s_ = *β*, for the *in cis design* we set *β*_ms_ = *β*. For a particular value of mean GFP steady-state, we can obtain lower coefficient of variation (meaning lower noise) in the *in cis* design in comparison to the *in trans* design and the ‘no sRNA’ case. For example, for an average 𝔼(GFP) = 10.4 molecules we have *β* = 0.1 [nM/min], η = 0.67 without the sRNA repression, for *k*_rep_ = 0.5 [1/(nM min)] we have 𝔼(GFP) = 9.29 molecules, η = 0.49, *β*= 0.3 [nM/min] in the *in cis* design and 𝔼(GFP) = 11.45 molecules, η = 0.69, *β*= 0.3 [nM/min] for the *in trans* design.

Additional simulations (see Figure S3 in SI) for the *in cis* design revealed that the level of noise attenuation can be tuned by several parameters: the repression strength *k*_rep_, the ribozyme cleavage rate *k*_rc_ and the degradation rate of sRNA *δ*_s_. In particular, increasing the ribozyme cleavage rate *k*_rc_ or the sRNA degradation rate *δ*_s_ lead to a decrease in the noise levels.

## 2.4 *In cis* module tunes the feedback strength and reduces noise in the TetR autorepressor

We then proceeded to investigate how the two modules behave in a feedback interconnection, such as for example when an *tetR-gfp* fusion gene is placed under the control of a P_tet_ promoter. In this case, we expect the TetR being produced to repress the activity of P_tet_ (Figure 6.I). We consider the following chemical reactions for the *in trans* design: 
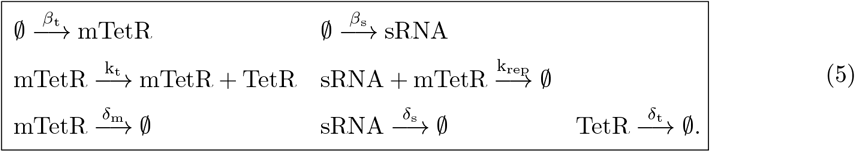

Here, GFP production is not modelled since it is fused to TetR and only serves as a reporter on TetR production. Both TetR and sRNA are controlled by *P*_tet_ (see SI for a full model description). We assume that the rest of the interactions follow mass action kinetics.

We assume that *β*_t_ = *β*_s_ to aid comparison with the *in cis* design, which can be modelled using the following chemical reactions: 
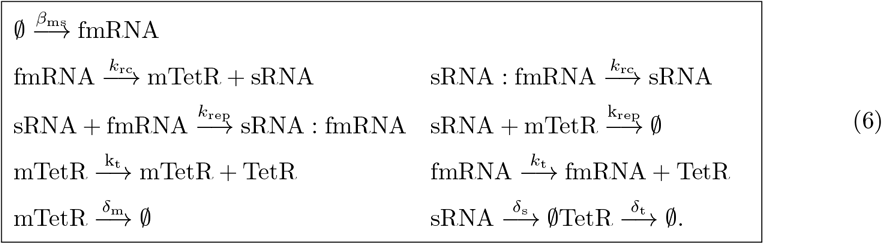

For the stochastic simulations in Figure 6.II we used the parameters/parameter ranges in Table S1 and additionally set *k*_rc_ = 1 [1/min], *k*_t_ = 1 [1/min]. These simulations suggest that the *in cis* design attenuates noise better in comparison with the no sRNA (classical autorepressor) circuit for a wider range of parameters than the *in trans* design. Note that with sufficient increase of the feedback strength the noise levels can be amplified by the *in trans* design, which is consistent with previous studies [Kelly et al., 2018]. In our *in cis* design the noise amplification does not occur for the simulated range of parameters (noise amplification is still possible for larger aTc concentrations). Furthermore, for a particular mean TetR level we can always select a combination of the sRNA repression strength k_rep_ and the aTc concentration so that the coefficient of variation is reduced in comparison to ‘no sRNA’ (see caption toFigure 6.II). In the *in trans* design these tuning dials are less effective: the noise reduction can be insignificant or the sRNA repression strength is very small, which means that the *in trans* approach *is not appropriate for noise reduction*. The noise analysis suggests that the repression rate *k*_rep_ adds a valuable tuning dial to the feedback strength design along with the aTc concentration. The numerical values of these simulations are given in Table S3 in SI.

**Figure 6.**
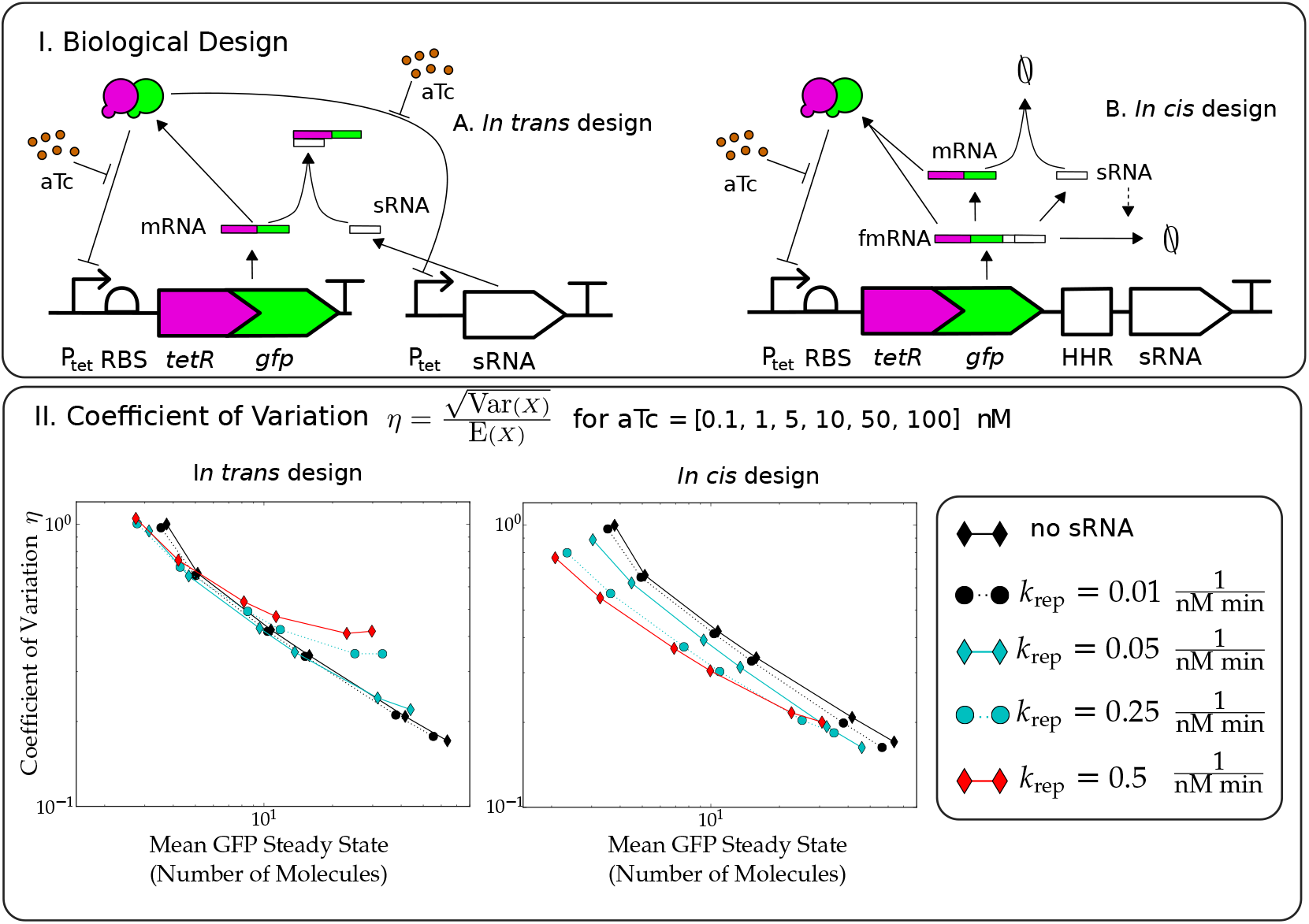
Improvement of the autorepressor design. I. Conceptual designs of the *in trans* and the *in cis* feedback strength regulators of the autorepressor. II. *In cis* filter effectively reduces noise in comparison to *in trans* filter, while both tune the feedback strength. We plot the coefficient of variation versus the mean GFP steady-state. Every data point corresponds to a different aTc concentration from [0.1, 1, 5, 10, 50, 100] [nM]. Every line corresponds to a different repression strength *k*_rep_. For a particular value of mean GFP steady-state, we can obtain a lower coefficient of variation (meaning lower noise) in the *in cis* design in comparison to the *in trans* design and the ‘no sRNA’ case, which corresponds to the classical autorepressor in both plots. For example, for an average 𝔼(TetR) = 10.7 molecules we have [aTc] = 5 [nM], η = 0.42 with the classical autorepressor (no sRNA), for *k*_rep_ = 0.25 [1/(nM min)] and [aTc] = 10 [nM] we have 𝔼(TetR) = 10.9 molecules, n = 0.30 in the *in cis* design and E(TetR) = 11.78 molecules, η = 0.42 in the *in trans* design.

## 3 Discussion

In this paper, we report the design of an sRNA-based feedback filter where the regulatory sRNA is placed directly after the gene to regulate *in cis* the signal, resulting in a filtered output. Modelling this new design, we showed that it can improve noise attenuation significantly compared to an in trans filter design and a no filter (no sRNA) design. Our results clearly indicate that the *in cis* design adapts better to the inputs than the *in trans* design mainly due to the presence of the feedback component. Moreover, in the *in cis* system, the production rate of the mRNA and sRNA change simultaneously, attenuating the transcription disturbance better than in the *in trans* design, where the relative gene expression rate varies significantly due to the sRNA and mRNA transcription rate being decoupled. Lastly, our *in cis* design requires less cellular resources (e.g. RNA polymerase), decreasing the burden imposed on the cell.

We successfully implemented this new sRNA-based filter *in vivo*. Our approach, using synthetic sRNA as described by [Na et al., 2013, Yoo et al., 2013] allows not only attenuation but also fine tuning to a desired output. Indeed, altering the length of the target binding sequence (TBS) allows varying the strength of the sRNA-mRNA binding, therefore leading to different levels of attenuation. We tested several length (from 22 to 30 nucleotides long) and could attenuate the output of the filter down to 40% of the unregulated output, very close to the values reported in other *in trans* designs [Kelly et al., 2018]. Increasing the length of the TBS should in theory allow higher attenuation levels, although off-target binding might then have to be taken in account [Na et al., 2013, Yoo et al., 2013]. Recently a similar architecture was proposed in mammalian cells [Lillacci et al., 2018], where micro RNA was placed *in cis* with the regulated gene. In our system, placing the syntethic sRNA *in cis* with the target mRNA without the ribozyme did not yield positive results. We showed, however, that the targeted mRNA and the regulatory sRNA have to be cleaved from each other for efficient output attenuation. Such cleavage was achieved by placing a self-cleaving hammerhead ribozyme (the HHR9 ribozyme) between the mRNA and the sRNA. The ribozyme represents another tuning dial allowing further fine tuning of the system. The ribozyme/synthetic sRNA approach, other than providing tuning dials such as the ribozyme cleavage rate and the repression strength, has another advantage: the repressing molecule is free from the active one, limiting possible unwanted effects. The emergence of synthetic ribozymes (self-cleaving or cleaving in response to a signal) should allow greater tuning flexibility.

Modelling both the *in cis* and *in trans* designs showed the clear advantages of the former design over the latter. While the mean steady-state behaviour of the two designs is quantitatively similar, the noise levels differ. In particular, the *in cis* design attenuates the transcription noise more efficiently thanks to the simultaneous bursts in transcription for the sRNA and the mRNA and the presence of feedback. Modelling suggests that the feedback strength in the filter is proportional to the ribozyme cleavage rate adding another benefit to the development of synthetic fast-cleaving ribozymes.

Further theoretical analysis showed that our design is a useful tool for feedback control design. We showed that the *in cis* design is well suited to tune down the feedback strength in a transcriptional based controller such as the TetR autorepressor. Again, the *in cis* design has superior noise properties in comparison to the *in trans* design. These findings are consistent with previously reported studies [Laurenti et al., 2018], where a Linear Noise Approximation [Van Kampen, 2007] was used to perform the noise attenuation analysis. Indeed, in a feedback setting, a given mean steady-state value can be achieved through either acting on the signal level (in our case aTc) or the strength of the feedback (in our case mRNA-sRNA binding): the *in cis* design offers a wide range of parameters achieving the same mean steady-state values with lower noise levels.

In this paper we presented a new sRNA-based feedback filter module. Together with the fast dynamics at which RNA operates, our in cis architecture is a simple, modular and tunable construct that can be applied in a wide range of synthetic biology applications while keeping the burden imposed on the cell at a minimum level.

## Author Contributions

ND and AS contributed equally to this work. Conceptualization, GHW and AP; Methodology, ND, AS, AP, GHW; Investigation, ND; Formal Analysis, AS; Writing – Original Draft, ND, AS; Writing – Review & Editing, ND, AS, AP, GHW; Funding Acquisition, AP; Resources, AP and GHW.

## Acknowledgements

The authors would like to thank Prof J.Keasling for providing a plasmid, Prof De la Peña for advising on the hammerhead ribozyme. This work is supported by UK’s Engineering and Physical Sciences (EPSRC) Grant EP/M002454/1.

## Declaration of Interests

The authors declare no competing interests.

## 4 Star Methods

### 4.1 Key resources table

| REAGENT OR RESOURCE | REFERENCE | IDENTIFIER |
| --- | --- | --- |
| CHEMICALS, PEPTIDES, AND RECOMBINANT PROTEINS |  |  |
| anhydrous tetracycline | TOKU-E | T055 |
| EXPERIMENTAL MODELS: ORGANISMS/STRAINS |  |  |
| *E. coli*: MG1655 | [Blattner et al., 1997] | ATCC47076 |
| RECOMBINANT DNA |  |  |
| Plasmid: pBbS2a-RFP | Shared by Prof J. Keasling [Lee et al., 2011] | addGene: 35328 |
| Plasmid: pND113 | This work | will be provided by GenBank |
| Plasmid: pND149 | This work | will be provided by GenBank |
| Plasmid: pND179 | This work | will be provided by GenBank |
| Plasmid: pND218 | This work | will be provided by GenBank |
| Plasmid: pND219 | This work | will be provided by GenBank |
| Plasmid: pND221 | This work | will be provided by GenBank |
| SOFTWARE AND ALGORITHMS |  |  |
| SnapGene | GSL Biotech LLC | http://www.snapgene.com |
| DINAMelt | [Markham and Zuker, 2005]<br>[Markham and Zuker, 2008] | http://mfold.rna.albany.edu/?q=DINAMelt/Two-state-melting |
| MATLAB R2016b | MathWorks, Inc., Natick, MA, USA | http://www.mathworks.com |
| cuda-sim | [Zhou et al., 2011] | https://github.com/jamesscottbrown/cuda-sim |

### 4.2 Contact for resource sharing

Further information and requests for resources should be addressed to Prof Papachristodoulou.

## 4.3 Method Details

### 4.3.1 Bacterial strains and plasmids

*Escherichia coli* MG1655 cells were used throughout this entire study unless stated otherwise. Plasmids were produced using standard cloning techniques. All synthetic DNA fragments (gBlocks) and primers used in this study were synthesized by Integrated DNA Technologies Inc. The length of the target binding sequences within the sRNA sequence were estimated using the web-based service DINAMelt (http://mfold.rna.albany.edu/?q=DINAMelt/Two-state-melting). We used pBbS2a-RFP (JBEI-2549, shared by Prof. J. Keasling) as a backbone for all the plasmids made for this work [Lee et al., 2011]. A list and a description of plasmids used in this study can be found in Table S4 in SI. Sequences of all plasmids have been submitted to GenBank. Full details are provided in SI.

### 4.3.2 Growth conditions and assays

Cells were grown overnight from single colonies to stationary phase in minimal medium 9 (M9) complemented with thiamine 0.34 mg/mL and ampicillin (100*μ*g/mL) at 30° C with shaking and then diluted 1/100 into fresh M9 with ampicillin (100*μ*g/mL) of cells were then loaded onto a 96-well plate (Corning) and left to grow for 2h at 30° C with shaking in a FLUOstar Omega Microplate Reader (BMG LABTECH). After this time, anhydrous tetracycline (aTc) at the appropriate concentration was added to the cells and measurements were acquired in the plate reader (gain: 1000). Absorbance and GFP fluorescence (excitation and emission wavelengths: 485 and 530 nm, with20 nm bandwidth, respectively) were measured every 3 minutes. Fluorescence was normalised by absorbance and plotted over time.

### 4.3.3 Mathematical modelling

We used mass action and Hill kinetic formalisms in order to model the chemical reactions as a Chemical Master Equation [Van Kampen, 2007]. The stochastic simulations were performed using the modified version (https://github.com/jamesscottbrown/cuda-sim) of the software tool cuda-sim [Zhou et al., 2011], which implements the Gillespie stochastic simulation algorithm. The deterministic simulations were performed in MATLAB using a built-in ordinary differential equation solver ODE15S. The parameter fitting was performed using non-linear least squares routine FIT in MATLAB.

## References

Agrawal, D., Tang, X., Westbrook, A., Marshall, R., Maxwell, C., Lucks, J., Noireaux, V., Beisel, C., Dunlop, M. and Franco, E. (2018). Mathematical modeling of RNA-based architectures for closed loop control of gene expression. ACS synthetic biology 7, 1219–1228.

Ang, J. and McMillen, D. (2013). Physical constraints on biological integral control design for homeostasis and sensory adaptation. Biophysical journal 104, 505–515.

Åstrom, K. and Murray, R. (2008). Feedback Systems: An Introduction for Scientists and Engineers. Princeton University Press, Princeton, NJ, USA.

Blattner, F., Plunkett, G., Bloch, C., Perna, N., Burland, V., Riley, M., Collado-Vides, J., Glasner, J., Rode, C., Mayhew, G. F. et al. (1997). The complete genome sequence of Escherichia coli K-12. Science 277, 1453–1462.

Briat, C., Gupta, A. and Khammash, M. (2016). Antithetic integral feedback ensures robust perfect adaptation in noisy biomolecular networks. Cell systems 2, 15–26.

Brophy, J. and Voigt, C. (2014). Principles of genetic circuit design. Nat methods 11, 508–520.

Cantone, I., Marucci, L., Iorio, F., Ricci, M., Belcastro, V., Bansal, M., Santini, S., Di Bernardo, M., Di Bernardo, D. and Cosma, M. (2009). A yeast synthetic network for in vivo assessment of reverse-engineering and modeling approaches. Cell 137, 172–181.

Cech, T. and Steitz, J. (2014). The noncoding RNA revolution-trashing old rules to forge new ones. Cell 157, 77–94.

Ceroni, F., Algar, R., Stan, G. and Ellis, T. (2015). Quantifying cellular capacity identifies gene expression designs with reduced burden. Nature methods 12, 415.

De la Peña, M., Gago, S. and Flores, R. (2003). Peripheral regions of natural hammerhead ribozymes greatly increase their self-cleavage activity. The EMBO journal 22, 5561–5570.

De La Peña, M. and García-Robles, I. (2010). Intronic hammerhead ribozymes are ultraconserved in the human genome. EMBO reports 11, 711–716.

Del Vecchio, D. and Murray, R. M. (2015). Biomolecular feedback systems. Princeton University Press Princeton, NJ.

Del Vecchio, D., Ninfa, A. J. and Sontag, E. D. (2008). Modular cell biology: retroactivity and insulation. Molecular systems biology 4, 161.

El-Samad, H., Goff, J. and Khammash, M. (2002). Calcium homeostasis and parturient hypocalcemia: an integral feedback perspective. Journal of theoretical biology 214, 17–29.

Folliard, T., Steel, H., Prescott, T., Wadhams, G., Rothschild, L. and Papachristodoulou, A. (2017). A synthetic recombinase-based feedback loop results in robust expression. ACS synthetic biology 6, 1663–1671.

Freemont, P. and Kitney, R. (2015). Synthetic Biology-a Primer (revised Edition). World Scientific.

Ghodasara, A. and Voigt, C. (2017). Balancing gene expression without library construction via a reusable sRNA pool. Nucleic acids research 45, 8116–8127.

Gottesman, S. and Storz, G. (2011). Bacterial small RNA regulators: versatile roles and rapidly evolving variations. Cold Spring Harbor perspectives in biology 3, a003798.

Haykin, S. (2002). Adaptive filter theory.

Holmqvist, E., Unoson, C., Reimegård, J. and Wagner, E. (2012). A mixed double negative feedback loop between the sRNA MicF and the global regulator Lrp. Molecular microbiology 84, 414–427.

Hooshangi, S., Thiberge, S. and Weiss, R. (2005). Ultrasensitivity and noise propagation in a synthetic transcriptional cascade. Proceedings of the National Academy of Sciences 102, 3581–3586.

Hsiao, V., De Los Santos, E., Whitaker, W., Dueber, J. and Murray, R. (2014). Design and implementation of a biomolecular concentration tracker. ACS synthetic biology 4, 150–161.

Hsiao, V., Swaminathan, A. and Murray, R. (2018). Control Theory for Synthetic Biology: Recent Advances in System Characterization, Control Design, and Controller Implementation for Synthetic Biology. IEEE Control Systems 38, 32–62.

Hu, C., Takahashi, M., Zhang, Y. and Lucks, J. (2018). Engineering a Functional small RNA Negative Autoregulation Network with Model-guided Design. ACS synthetic biology 7, 1507–1518.

Hussein, R. and Lim, H. (2012). Direct comparison of small RNA and transcription factor signaling. Nucleic acids research 40, 7269–7279.

Iglesias, P. and Ingalls, B. (2010). Control theory and systems biology. MIT Press.

Kelly, C., Harris, A., Steel, H., Hancock, E., Heap, J. and Papachristodoulou, A. (2018). Synthetic negative feedback circuits using engineered small RNAs. Nucleic acids research 46, 9875–9889.

Laurenti, L., Csikasz-Nagy, A., Kwiatkowska, M. and Cardelli, L. (2018). Molecular Filters for Noise Reduction. Biophysical Journal 114, 3000–3011.

Lee, T., Krupa, R., Zhang, F., Hajimorad, M., Holtz, W., Prasad, N., Lee, S. and Keasling, J. (2011). BglBrick vectors and datasheets: a synthetic biology platform for gene expression. Journal of biological engineering 5, 12.

Lillacci, G., Benenson, Y. and Khammash, M. (2018). Synthetic control systems for high performance gene expression in mammalian cells. Nucleic acids research 46, 9855–9863.

Liu, X., Zhou, P. and Wang, R. (2013). Small RNA-mediated switch-like regulation in bacterial quorum sensing. IET systems biology 7, 182–187.

Livny, J. and Waldor, M. (2007). Identification of small RNAs in diverse bacterial species. Current opinion in microbiology 10, 96–101.

Markham, N. and Zuker, M. (2005). DINAMelt web server for nucleic acid melting prediction. Nucleic acids research 33, W577-W581.

Markham, N. and Zuker, M. (2008). UNAFold. In Bioinformatics pp. 3–31. Springer.

Mehta, P., Goyal, S. and Wingreen, N. (2008). A quantitative comparison of sRNA-based and protein-based gene regulation. Molecular systems biology 4, 221.

Menolascina, F., Di Bernardo, M. and Di Bernardo, D. (2011). Analysis, design and implementation of a novel scheme for in-vivo control of synthetic gene regulatory networks. Automatica, Special Issue on Systems Biology 47, 1265–1270.

Michaux, C., Verneuil, N., Hartke, A. and Giard, J.-C. (2014). Physiological roles of small RNA molecules. Microbiology 160, 1007–1019.

Milias-Argeitis, A., Summers, S., Stewart-Ornstein, J., Zuleta, I., Pincus, D., El-Samad, H., Khammash, M. and Lygeros, J. (2011). In silico feedback for in vivo regulation of a gene expression circuit. Nat biotechnol 29, 1114–1116.

Muranaka, N. and Yokobayashi, Y. (2010). A synthetic riboswitch with chemical band-pass response. Chemical communications 46, 6825–6827.

Na, D., Yoo, S., Chung, H., Park, H., Park, J. and Lee, S. (2013). Metabolic engineering of Escherichia coli using synthetic small regulatory RNAs. Nature biotechnology 31, 170.

Nitzan, M., Rehani, R. and Margalit, H. (2017). Integration of bacterial small RNAs in regulatory networks. Annual review of biophysics 46, 131–148.

Ozdemir, T., Fedorec, A., Danino, T. and Barnes, C. (2018). Synthetic Biology and Engineered Live Biotherapeutics: Toward Increasing System Complexity. Cell systems 7, 5–16.

Perreault, J., Weinberg, Z., Roth, A., Popescu, O., Chartrand, P., Fer-beyre, G. and Breaker, R. (2011). Identification of hammerhead ribozymes in all domains of life reveals novel structural variations. PLoS computational biology 7, e1002031.

Purnick, P. and Weiss, R. (2009). The second wave of synthetic biology: from modules to systems. Nat. Rev. Mol. Cell Biol. 10, 410–422.

Robledo, M., Schlüter, J., Linne, U., Albaum, S., Jimenéz-Zurdo, J., Becker, A. et al. (2018). An sRNA and cold shock protein homolog-based feedforward loop post-transcriptionally controls cell cycle master regulator CtrA. Frontiers in microbiology 9, 763.

Rosenfeld, N., Elowitz, M. and Alon, U. (2002). Negative autoregulation speeds the response times of transcription networks. Journal of molecular biology 323, 785–793.

Samoilov, M., Arkin, A. and Ross, J. (2002). Signal processing by simple chemical systems. The Journal of Physical Chemistry A 106, 10205–10221.

Sohka, T., Heins, R., Phelan, R., Greisler, J., Townsend, C. and Ostermeier, M. (2009). An externally tunable bacterial band-pass filter. Proc National Acad Sciences 106, 10135–10140.

Steel, H., Harris, A., Hancock, E., Kelly, C. and Papachristodoulou, A. (2017a). Frequency domain analysis of small non-coding RNAs shows summing junction-like behaviour. In Proc Conf Decision Control pp. 5328–5333, IEEE.

Steel, H., Lillacci, G., Khammash, M. and Papachristodoulou, A. (2017b). Challenges at the interface of control engineering and synthetic biology. In Proceedings of the IEE Conference on Decision and Control pp. 1014–1023, IEEE.

Takahashi, M., Chappell, J., Hayes, C., Sun, Z., Kim, J., Singhal, V., Spring, K., Al-Khabouri, S., Fall, C., Noireaux, V. et al. (2014). Rapidly characterizing the fast dynamics of RNA genetic circuitry with cell-free transcription-translation (TX-TL) systems. ACS synthetic biology 4, 503–515.

Thattai, M. and van Oudenaarden, A. (2002). Attenuation of noise in ultrasensitive signaling cascades. Biophysical journal 82, 2943–2950.

Uhlendorf, J., Miermont, A., Delaveau, T., Charvin, G., Fages, F., Bottani, S., Batt, G. and Hersen, P. (2012). Long-term model predictive control of gene expression at the population and single-cell levels. Proc. Nat. Academy Sciences 109, 14271–14276.

Van Kampen, N. (2007). Stochastic Processes in Physics and Chemistry. Elsevier.

Yoo, S., Na, D. and Lee, S. (2013). Design and use of synthetic regulatory small RNAs to control gene expression in Escherichia coli. Nature protocols 8, 1694.

Zechner, C., Seelig, G., Rullan, M. and Khammash, M. (2016). Molecular circuits for dynamic noise filtering. Proc National Acad Sciences 113, 4729–4734.

Zhou, S. (2016). Synthetic biology: bacteria synchronized for drug delivery. Nature 536, 33.

Zhou, Y., Liepe, J., Sheng, X., Stumpf, M. and Barnes, C. (2011). GPU accelerated biochemical network simulation. Bioinformatics 27, 874–876.

