## Supplementary Information for "Design of a Synthetic sRNA-based Feedback Filter Module"

#### S1 Modelling of the sRNA filters

##### S1.1 Modelling of the *in trans* design

First, we briefly discuss an *in trans* filter, which we model by means of the following chemical reaction network,

$$\boxed{\begin{array}{lll} \emptyset \xrightarrow{\beta_{\mathrm{m}}} \mathrm{mGFP} & \emptyset \xrightarrow{\beta_{\mathrm{s}}} \mathrm{sRNA} & \\ \mathrm{sRNA} + \mathrm{mGFP} \xrightarrow{k_{\mathrm{rep}}} \emptyset & \mathrm{mGFP} \xrightarrow{k_{\mathrm{t}}} \mathrm{mGFP} + \mathrm{GFP} & \\ \mathrm{mGFP} \xrightarrow{\delta_{\mathrm{m}}} \emptyset & \mathrm{sRNA} \xrightarrow{\delta_{\mathrm{s}}} \emptyset & \mathrm{GFP} \xrightarrow{\delta_{\mathrm{p}}} \emptyset, \end{array}} \tag{S1}$$

where mGFP (the mRNA of the GFP gene) and sRNA are transcribed at rates $\beta_{\mathrm{m}}$ and $\beta_{\mathrm{s}}$, respectively. The sRNA binds to mGFP with rate $k_{\mathrm{rep}}$, and mGFP is translated to GFP at a rate $k_t$. We assume that the degradation/dilution rates for all the species are different, since it is known that mRNAs generally degrade faster than proteins, and the reported values of the degradation rate of sRNA vary [Hussein and Lim, 2012]. It is assumed that the Hfq protein can stabilise the sRNA's, however, there is evidence that this stabilisation occurs after sRNA and mRNA binding [Hussein and Lim, 2012], and the sRNA degradation rate typically varies between $\delta_{\mathrm{m}}$ and $\delta_{\mathrm{p}}$. We do not model unbinding of sRNA from the target mRNA by assuming that the rate of this reaction is negligibly small. This modelling assumption was used in the literature before [Hussein and Lim, 2012, Steel et al., 2017, Kelly et al., 2018, Agrawal et al., 2018]. Assuming mass action kinetics for all reactions, we obtain the following Ordinary Differential Equation (ODE) Model:

$$\boxed{\begin{aligned} \frac{d}{dt}[\mathrm{mGFP}] &= \beta_{\mathrm{m}} - \delta_{\mathrm{m}}[\mathrm{mGFP}] - k_{\mathrm{rep}}[\mathrm{sRNA}][\mathrm{mGFP}], \\ \frac{d}{dt}[\mathrm{sRNA}] &= \beta_{\mathrm{s}} - \delta_{\mathrm{s}}[\mathrm{sRNA}] - k_{\mathrm{rep}}[\mathrm{sRNA}][\mathrm{mGFP}], \\ \frac{d}{dt}[\mathrm{GFP}] &= k_{\mathrm{t}}[\mathrm{mGFP}] - \delta_{\mathrm{p}}[\mathrm{GFP}], \end{aligned}} \tag{S2}$$

where $[X]$ denotes the concentration of species $X$. For the sake of comparison to the *in cis* design, we assume that the strengths of both promoters in model (S2) are constant and tunable. For example, the promoters are inducible and we can vary the concentrations of the inducers.

##### S1.2 Detailed modelling of the *in cis* design

For the *in cis* design, we assume that the relevant gene sequence after the promoter and ribosome binding site is as follows: mGFP, ribozyme, and sRNA. A self-cleaving ribozyme [Weinberg et al., 2015], [Hammann et al., 2012], [Perreault et al., 2011] is a ribonucleic acid enzyme, which cleaves the functional mRNA into two parts, which in our case are the mRNA of the target gene and the sRNA. After cleavage the sRNA can bind to a specific region of mGFP, and thus silence its translation. We note that only free sRNA (unbound from the mRNA of the GFP) can bind to the mRNAs, as experimental data suggests. Since the functional mRNA (mGFP-ribozyme-sRNA) also has this region, we assume that the free sRNA can bind to the functional mRNA as well. The resulting complex of the sRNA and the functional mRNA can also be cleaved by a ribozyme resulting in another sRNA and the mRNA-sRNA complex which is however inactive and we do not consider. Both the mRNA of the target gene and the functional mRNA can be translated into the target protein. We assume that the functional mRNA fmRNA is transcribed with rate $\beta_{\mathrm{ms}}$. We assume that the self-cleaving of the ribozyme takes place after the transcription process. Hence, fmRNA is cleaved to mGFP, and sRNA with a rate $k_{\mathrm{rc}}$. In total, we have the following chemical reaction model:

$$\boxed{\begin{array}{ll} \emptyset \xrightarrow{\beta_{\mathrm{ms}}} \mathrm{fmRNA} & \\ \mathrm{fmRNA} \xrightarrow{k_{\mathrm{rc}}} \mathrm{mGFP} + \mathrm{sRNA} & \mathrm{sRNA} : \mathrm{fmRNA} \xrightarrow{k_{\mathrm{rc}}} \mathrm{sRNA} \\ \mathrm{sRNA} + \mathrm{fmRNA} \xrightarrow{k_{\mathrm{rep}}} \mathrm{sRNA} : \mathrm{fmRNA} & \mathrm{sRNA} + \mathrm{mGFP} \xrightarrow{k_{\mathrm{rep}}} \emptyset \\ \mathrm{mGFP} \xrightarrow{k_{\mathrm{t}}} \mathrm{mGFP} + \mathrm{GFP} & \mathrm{fmRNA} \xrightarrow{k_{\mathrm{t}}} \mathrm{fmRNA} + \mathrm{GFP} \\ \mathrm{mGFP} \xrightarrow{\delta_{\mathrm{m}}} \emptyset,\ \mathrm{sRNA} \xrightarrow{\delta_{\mathrm{s}}} \emptyset,\ \mathrm{GFP} \xrightarrow{\delta_{\mathrm{p}}} \emptyset. & \end{array}} \tag{S3}$$

Using the mass-action kinetics formalism we obtain the following differential equations:

$$\begin{aligned} \frac{d}{dt}[\mathrm{mGFP}] &= k_{\mathrm{rc}}[\mathrm{fmRNA}] - \delta_{\mathrm{m}}[\mathrm{mGFP}] - k_{\mathrm{rep}}[\mathrm{sRNA}][\mathrm{mGFP}], \\ \frac{d}{dt}[\mathrm{GFP}] &= k_{\mathrm{t}}([\mathrm{mGFP}] + [\mathrm{fmRNA}]) - \delta_{\mathrm{p}}[\mathrm{GFP}], \\ \frac{d}{dt}[\mathrm{sRNA}] &= k_{\mathrm{rc}}([\mathrm{sRNA} : \mathrm{fmRNA}] + [\mathrm{fmRNA}]) - \delta_{\mathrm{s}}[\mathrm{sRNA}] - \\ &\quad - k_{\mathrm{rep}}[\mathrm{sRNA}]([\mathrm{mGFP}] + [\mathrm{fmRNA}]), \\ \frac{d}{dt}[\mathrm{fmRNA}] &= \beta_{\mathrm{ms}} - \delta_{\mathrm{m}}[\mathrm{fmRNA}] - k_{\mathrm{rc}}[\mathrm{fmRNA}] - \\ &\quad - k_{\mathrm{rep}}[\mathrm{sRNA}][\mathrm{fmRNA}], \\ \frac{d}{dt}[\mathrm{sRNA} : \mathrm{fmRNA}] &= k_{\mathrm{rep}}[\mathrm{sRNA}][\mathrm{fmRNA}] - \delta_{\mathrm{s}}[\mathrm{sRNA} : \mathrm{fmRNA}] - \\ &\quad - k_{\mathrm{rc}}[\mathrm{sRNA} : \mathrm{fmRNA}]. \end{aligned} \tag{S4}$$

In the rest of this subsection, we describe the model simplification procedure. Let us introduce a new quantity $[\mathrm{mGFP}^{\mathrm{tot}}] = [\mathrm{mGFP}] + [\mathrm{fmRNA}]$, which allows us to eliminate the equations for mGFP, resulting in the model:

$$\begin{aligned} \frac{d}{dt}[\mathrm{mGFP}^{\mathrm{tot}}] &= \beta_{\mathrm{ms}} - \delta_{\mathrm{m}}[\mathrm{mGFP}^{\mathrm{tot}}] - k_{\mathrm{rep}}[\mathrm{sRNA}][\mathrm{mGFP}^{\mathrm{tot}}], \\ \frac{d}{dt}[\mathrm{GFP}] &= k_{\mathrm{t}}[\mathrm{mGFP}^{\mathrm{tot}}] - \delta_{\mathrm{p}}[\mathrm{GFP}], \\ \frac{d}{dt}[\mathrm{sRNA}] &= k_{\mathrm{rc}}([\mathrm{fmRNA}] + [\mathrm{sRNA} : \mathrm{fmRNA}]) - \delta_{\mathrm{s}}[\mathrm{sRNA}] - \\ &\quad - k_{\mathrm{rep}}[\mathrm{sRNA}][\mathrm{mGFP}^{\mathrm{tot}}] \\ \frac{d}{dt}[\mathrm{fmRNA}] &= \beta_{\mathrm{ms}} - \delta_{\mathrm{m}}[\mathrm{fmRNA}] - k_{\mathrm{rc}}[\mathrm{fmRNA}] - \\ &\quad - k_{\mathrm{rep}}[\mathrm{sRNA}][\mathrm{fmRNA}] \\ \frac{d}{dt}[\mathrm{sRNA} : \mathrm{fmRNA}] &= k_{\mathrm{rep}}[\mathrm{sRNA}][\mathrm{fmRNA}] - \delta_{\mathrm{s}}[\mathrm{sRNA} : \mathrm{fmRNA}] - \\ &\quad - k_{\mathrm{rc}}[\mathrm{sRNA} : \mathrm{fmRNA}]. \end{aligned} \tag{S5}$$

If $k_{\mathrm{rc}} \gg \delta_{\mathrm{p}}, \delta_{\mathrm{p}}, \delta_{\mathrm{p}}$ the species [fmRNA] and [sRNA : fmRNA] can be eliminated due to time-scale separation as we discuss in detail in what follows. In simple terms, the sum of the variables [fmRNA] and [sRNA : fmRNA] can be replaced by their values at the steady-state:

$$[\mathrm{fmRNA}] + [\mathrm{sRNA} : \mathrm{fmRNA}] = \frac{\beta_{\mathrm{ms}}}{\delta_{\mathrm{s}} + k_{\mathrm{rc}}} \frac{\delta_{\mathrm{s}} + k_{\mathrm{rc}} + k_{\mathrm{rep}}[\mathrm{sRNA}]}{\delta_{\mathrm{m}} + k_{\mathrm{rc}} + k_{\mathrm{rep}}[\mathrm{sRNA}]} = \frac{\beta_{\mathrm{ms}}}{\delta_{\mathrm{m}} + k_{\mathrm{rc}}} \frac{1 + \frac{k_{\mathrm{rep}}[\mathrm{sRNA}]}{\delta_{\mathrm{s}} + k_{\mathrm{rc}}}}{1 + \frac{k_{\mathrm{rep}}[\mathrm{sRNA}]}{\delta_{\mathrm{m}} + k_{\mathrm{rc}}}} \approx \frac{\beta_{\mathrm{ms}}}{k_{\mathrm{rc}} + \delta_{\mathrm{m}}},$$

where we assume that $1 + \frac{k_{\mathrm{rep}}[\mathrm{sRNA}]}{\delta_{\mathrm{s}} + k_{\mathrm{rc}}} \approx 1 + \frac{k_{\mathrm{rep}}[\mathrm{sRNA}]}{\delta_{\mathrm{m}} + k_{\mathrm{rc}}}$. We thus obtain the following model:

$$\boxed{\begin{aligned} \frac{d}{dt}[\mathrm{mGFP}^{\mathrm{tot}}] &= \beta_{\mathrm{ms}} - \delta_{\mathrm{m}}[\mathrm{mGFP}^{\mathrm{tot}}] - k_{\mathrm{rep}}[\mathrm{sRNA}][\mathrm{mGFP}^{\mathrm{tot}}], \\ \frac{d}{dt}[\mathrm{GFP}] &= k_{\mathrm{t}}[\mathrm{mGFP}^{\mathrm{tot}}] - \delta_{\mathrm{p}}[\mathrm{GFP}], \\ \frac{d}{dt}[\mathrm{sRNA}] &= \frac{k_{\mathrm{rc}}}{k_{\mathrm{rc}} + \delta_{\mathrm{m}}}\beta_{\mathrm{ms}} - \delta_{\mathrm{s}}[\mathrm{sRNA}] - k_{\mathrm{rep}}[\mathrm{sRNA}][\mathrm{mGFP}^{\mathrm{tot}}]. \end{aligned}} \tag{S6}$$

This analysis suggests that the ribozymes (and their cleavage rate $k_{\mathrm{rc}}$) can be used to adjust the gain of the attenuation, as well as the sRNA-mGFP binding strength $k_{\mathrm{rep}}$. With the ribozyme cleavage rate increasing the *in cis* deterministic model converges to the *in trans* deterministic model subject to adjusting the production rates for the sRNA and mRNA of the GFP, that is, setting $\beta_{\mathrm{m}} = \beta_{\mathrm{ms}}$, and $\beta_{\mathrm{s}} = \frac{k_{\mathrm{rc}}}{k_{\mathrm{rc}} + \delta_{\mathrm{m}}}\beta_{\mathrm{ms}}$. This indicates that under identical conditions the deterministic models of both designs (*in cis* and *in trans*) behave similarly provided that the ribozyme cleavage occurs on a much faster time-scale than the other reactions. However, there is a substantial difference in their response to changes in $\beta_m$, $\beta_s$ and $\beta_{ms}$: noise in the GFP promoter can be attenuated in the *in cis* system, while this is harder to achieve in the *in trans* system, which we illustrate in what follows. Additionally, the *in trans* system needs more cellular resources (e.g. RNA polymerase), and tuning the sRNA production rate requires tuning the promoter, whereas in the *in cis* system one can tune the ribozyme cleavage rate.

In order to fit the parameter values, we also compute the steady-state values of the simplified model (S6). A detailed derivation can be found in what follows, while here we present the resulting formula:

$$\begin{aligned} [\mathrm{GFP}]^{\mathrm{ss}} &= \frac{k_{\mathrm{t}}}{\delta_{\mathrm{p}}} \frac{\beta_{\mathrm{ms}}}{\delta_{\mathrm{m}} + k_{\mathrm{rep}}[\mathrm{sRNA}]^{ss}}, \\ [\mathrm{sRNA}]^{ss} &= -\frac{(\delta_{\mathrm{s}}\delta_{\mathrm{m}} + (1-\alpha)\beta_{\mathrm{ms}}k_{\mathrm{rep}})}{2k_{\mathrm{rep}}\delta_{\mathrm{s}}} + \\ &\qquad \frac{\sqrt{(\delta_{\mathrm{s}}\delta_{\mathrm{m}} + (1-\alpha)\beta_{\mathrm{ms}}k_{\mathrm{rep}})^2 + 4k_{\mathrm{rep}}\alpha\beta_{\mathrm{ms}}\delta_{\mathrm{m}}\delta_{\mathrm{s}}}}{2k_{\mathrm{rep}}\delta_{\mathrm{s}}}, \end{aligned}$$

where $\alpha = \frac{k_{\mathrm{rc}}}{k_{\mathrm{rc}} + \delta_{\mathrm{m}}}$. If we assume that the ribozyme cleavage rate $k_{\mathrm{rc}}$ is substantially large (that is $\alpha \approx 1$), and $\frac{k_{\mathrm{rep}}\beta_{\mathrm{ms}}}{\delta_{\mathrm{m}}\delta_{\mathrm{s}}}$ is much larger than one we have

$$[\mathrm{GFP}]^{\mathrm{ss}} = \frac{k_{\mathrm{t}}}{\delta_{\mathrm{p}}} \frac{2\beta_{\mathrm{ms}}/\delta_{\mathrm{m}}}{1 + \sqrt{1 + 4\frac{k_{\mathrm{rep}}\beta_{\mathrm{ms}}}{\delta_{\mathrm{m}}\delta_{\mathrm{s}}}}} \approx \frac{k_{\mathrm{t}}}{\delta_{\mathrm{p}}}\sqrt{\frac{\beta_{\mathrm{ms}}\delta_{\mathrm{s}}}{k_{\mathrm{rep}}\delta_{\mathrm{m}}}}.$$

This approximation indicates that the ratio between the output (GFP concentration) and the input ($\beta_{\mathrm{ms}}$) can be estimated for a wide parameter range without taking the values of other parameters into account.

#### S1.3 Parameter values and fitting

Some parameter values can be taken form the literature. For example, we make a fairly standard assumption that the culture doubling time is 30 [min] [Liang et al., 1999], and hence the dilution rate is 0.0234 [1/min], which we set as the dilution rate for the proteins. We set the half-life of the mRNA to 2.8 [min], hence the degradation rate is 0.2476 [1/min] [Chen et al., 2015, McCullen et al., 2010]. The half-life of the sRNA can vary according to the previous studies [Hussein and Lim, 2012]. It was suggested that because of the sRNA stabilisation by the Hfq protein the sRNA degradation rate is typically an order of magnitude lower than the degradation rate of mRNA. Therefore we vary the dilution/degradation rate of the sRNA between 0.0482 (summing up the protein dilution and the sRNA degradation rates as in [Kelly et al., 2018]) and 0.2476 [1/min] (the mRNA degradation rate). We vary the ribozyme cleavage rate between 1 and 5 [1/min] as was reported in [Weinberg et al., 2015, Hammann et al., 2012, Perreault et al., 2011] for different ribozymes. Finally, we set the translation rate to $k_t = 1$ [1/min], which lies within reasonable biological bounds [Hussein and Lim, 2012].

In order to fit the parameters we will use the steady-state data and perform the fit in two stages: First, we identify the parameters of mRNA production, and then we identify the repression strengths. In order to identify the promoter parameters, we consider the case with no sRNA in the loop. In particular, we use the model for the $P_{\text{tet}}$ promoter [Tamsir et al., 2011] and in this case we have:

Table S1: Parameter values for theoretical analysis

| parameter | value/interval | units |
|---|---|---|
| $\delta_{\text{p}}$ | 0.0234 | 1/min |
| $\delta_{\text{m}}$ | 0.2476 | 1/min |
| $\delta_{\text{s}}$ | [0.0482, 0.2476] | 1/min |
| $k_{\text{rc}}$ | [1, 5] | 1/min |
| $k_{\text{t}}$ | 1 | 1/min |
| $\beta_{\text{ms}}$ | [0, 1] | nM/min |
| $k_{\text{rep}}$ | [0, 0.5] | 1/(nM min) |

$$Y_0 = \frac{K_0}{1 + K_1 + 2\dfrac{K_2}{1 + X/K_D} + \left(\dfrac{K_2}{1 + X/K_D}\right)^2}, \quad K_2 = \tilde{K}_2[\text{TetR}]$$

where $X$ is the concentration of anhydrotetracycline (aTc), $Y_0$ is the normalised (with respect to the maximum value) fluorescence in the construct with no sRNA, and [TetR] is the concentration of the free TetR. Naturally, we cannot identify all the parameters separately and hence we compute $K_0$, $K_1$, $K_2$, and $K_D$. After that we use the following form in order to fit $k_{\text{rep}}$:

$$Y = \frac{2Y_0}{1 + S}$$
$$S = \frac{-(1-\alpha)Y_0 k_{\text{rep}}}{\delta_{\text{m}}\delta_{\text{s}}} + \sqrt{\left(1 + \frac{(1-\alpha)Y_0 k_{\text{rep}}}{\delta_{\text{m}}\delta_{\text{s}}}\right)^2 + 4\frac{k_{\text{rep}}\alpha Y_0}{\delta_{\text{m}}\delta_{\text{s}}}},$$

In order to fit the parameters we use nonlinear least squares methods implemented in MATLAB. We obtain the following parameter values for the promoter activity:

$$K_0 = 2.26\,[\text{nM/min}], \quad K_1 = 1.054, \quad K_2 = 18.46, \quad K_D = 0.1182\,[\text{nM}],$$

where the variables $K_1$ and $K_2$ are dimensionless. These values are generally consistent with those reported in the literature [Kennell and Riezman, 1977] and [Liang et al., 1999].

We fit the parameter $k_{\text{rep}}$ also using nonlinear least squares methods for different sRNA degradation rates in order to estimate a possible range for the binding rate and obtain the following parameters:

$$k_{\rm rc} = 5\,[1/\text{min}],\quad \delta_{\rm s} = 0.0482\,[1/\text{min}],\quad k_{\rm rep} = \begin{pmatrix}0.0080 & 0.0173 & 0.0438\end{pmatrix}\,[1/(\text{nM min})],$$
$$k_{\rm rc} = 5\,[1/\text{min}],\quad \delta_{\rm s} = 0.1\,[1/\text{min}],\quad k_{\rm rep} = \begin{pmatrix}0.0166 & 0.0359 & 0.0909\end{pmatrix}\,[1/(\text{nM min})],$$
$$k_{\rm rc} = 5\,[1/\text{min}],\quad \delta_{\rm s} = 0.2476\,[1/\text{min}],\quad k_{\rm rep} = \begin{pmatrix}0.0412 & 0.0888 & 0.2251\end{pmatrix}\,[1/(\text{nM min})],$$
$$k_{\rm rc} = 1\,[1/\text{min}],\quad \delta_{\rm s} = 0.0482\,[1/\text{min}],\quad k_{\rm rep} = \begin{pmatrix}0.0103 & 0.0238 & 0.0698\end{pmatrix}\,[1/(\text{nM min})],$$
$$k_{\rm rc} = 1\,[1/\text{min}],\quad \delta_{\rm s} = 0.1\,[1/\text{min}],\quad k_{\rm rep} = \begin{pmatrix}0.0214 & 0.0494 & 0.1449\end{pmatrix}\,[1/(\text{nM min})],$$
$$k_{\rm rc} = 1\,[1/\text{min}],\quad \delta_{\rm s} = 0.2476\,[1/\text{min}],\quad k_{\rm rep} = \begin{pmatrix}0.0530 & 0.1223 & 0.3588\end{pmatrix}\,[1/(\text{nM min})],$$

where the repression rates $k_{\rm rep} = \begin{pmatrix}k^1_{\rm rep} & k^2_{\rm rep} & k^3_{\rm rep}\end{pmatrix}$ correspond to the number of base pairs $l = \begin{pmatrix}22 & 27 & 30\end{pmatrix}$.

As the reader can observe, we cannot distinguish between the parameters $\delta_{\rm s}$, $k_{\rm rep}$ based on our fitting formula, in fact one can verify that the ratio $k_{\rm rep}/\delta_{\rm s}$ remains roughly the same for different values of parameters with a fixed value of $k_{\rm rc}$. This means that we cannot identify simultaneously $\delta_{\rm s}$ and $k_{\rm rep}$ and we resorted to a value of $\delta_{\rm s}$ from the literature and the fitted value of $k_{\rm rep}$. Given these estimations we take the parameter values and ranges as presented in Table S1.

#### S1.4 Comparison of the models for *in cis* and *in trans* designs

In order to perform a fair comparison between the models we need to choose appropriate production rates for mRNA and sRNA for the two designs. We assume that all promoters in both designs are identical, and we set $\beta_{\rm m} = \beta_{\rm ms}$ and $\beta_{\rm s} = \beta_{\rm ms}$. Therefore, throughout the section we will vary the parameters $\beta_{\rm ms}$, $k_{\rm rep}$ and $\delta_{\rm s}$ in our simulations.

##### S1.4.1 Trajectories

We perform stochastic simulations using the direct Gillespie algorithm [Gillespie, 1977] implemented in the software tool CUDA-SIM [Zhou et al., 2011]. This software tool is written in C++ and uses the GPU, which significantly accelerates computations. First, we compute 500 trajec-

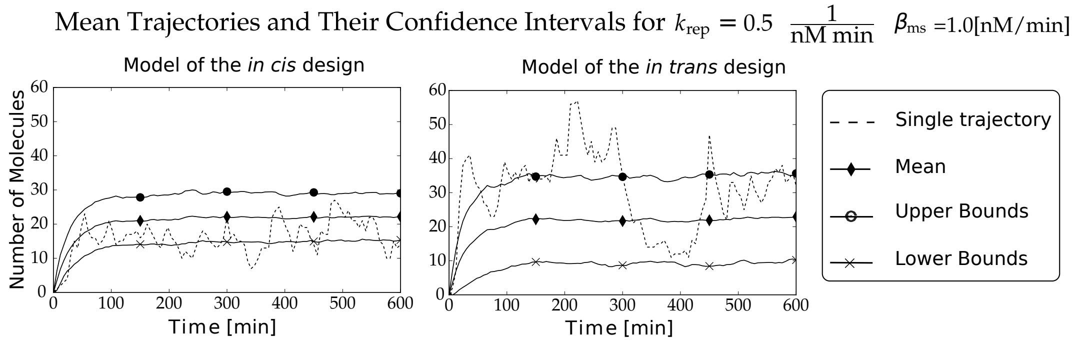


Figure S1: Mean GFP expression rates vs time for different values of parameters. We set $k_{\rm rep} = 0.5$ [1/(nM·min)], $\delta_{\rm s} = 0.0482$ [1/min], $k_{\rm rc} = 1$ [1/min], and the rest of the parameters as in Table S1. For the *in cis* system we set $\beta_{\rm tx-mrna} = 1$ [nM/min] and for the *in trans* system we set $\beta_{\rm m} = \beta_{\rm s} = 1$ [nM/min]. This figure shows that the transcriptional bursts in the *in cis* design can be attenuated by having sRNA, while in the *in trans* design these bursts will be much larger.

tories of the models on the time interval $[0, 600]$ [min] in order to evaluate the time evolution of the models. For the *in cis* design we set $k_{\rm rep} = 0.5$ [1/(nM·min)], $d_{\rm s} = 0.0482$ [1/min], $k_{\rm rc} = 1$ [1/min], and $\beta_{\rm ms} = 0.3$ and 1 [nM/min]. For the *in trans* design we compute the equivalent transcription rates and set $\beta_{\rm m} = \beta_{\rm s} = 0.3$ and 1 [nM/min]. The rest of the parameters are chosen as in Table S1. In Figure S1, we plot the results: The mean trajectories of both models converge fast to the mean steady-state values and the steady-state values are not significantly different between the two models. However, the trajectories of the *in trans* design appear to be noisier than the trajectories of the *in cis* design as there are significant variations in the individual trajectories and the confidence intervals (which we simply set to plus and minus standard deviations from the current value). We attribute the noisy behaviour to transcription bursts in mRNA and sRNA and the lack of feedback in the *in cis* design. In the *in trans* design the uncorrelated bursts in the production of mRNA and sRNA cause the sudden increase and the sudden decrease in the GFP production. The bursts in the production of mRNA and sRNA occur independently of each other due to different promoters. In the *in cis* design, these bursts occur simultaneously and some mRNA's are silenced by the sRNA's.

##### S1.4.2 Steady-state models

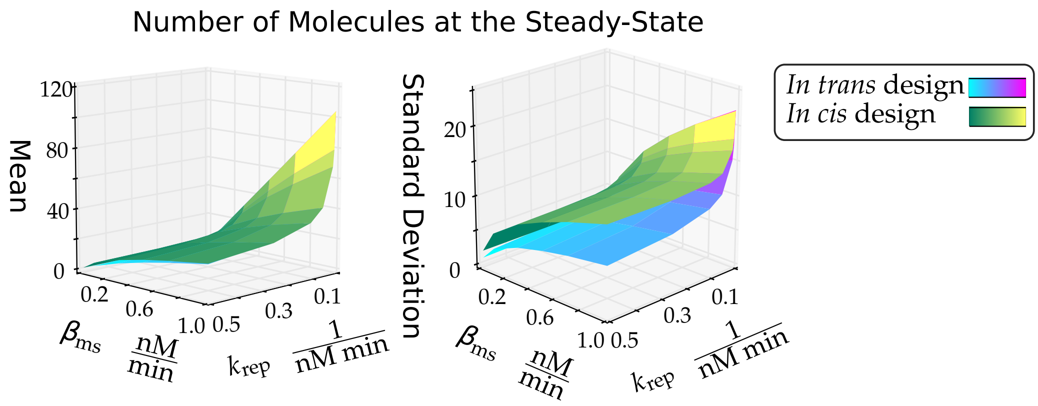


Figure S2: Model comparison for the *in trans* design and the *in cis* design systems. Statistics of GFP molecule number at $t = 2000$ [min] while performing a sweep over $\beta_{\text{ms}}$ and $k_{\text{rep}}$. We set $\beta_{\text{m}} = \beta_{\text{ms}}$, $\beta_{\text{s}} = \beta_{\text{ms}}$, $\delta_{\text{s}} = 0.0482$ [1/min], $k_{\text{rc}} = 1$, the rest of the parameters are as in Table S1. The mean number of molecules are similar for the two designs, while the standard deviations differ significantly.

We compute 50000 trajectories on the time interval $[0, 2000]$ [min]. At the time $t = 2000$ [min], we compute the means $\mathbb{E}[\text{GFP}(t)]$ and the standard deviations $\sqrt{\text{Var}[\text{GFP}(t)]}$. The results of the parameter sweeps over $\beta_{\text{ms}}$ and $k_{\text{rep}}$ are reported in Figure S2. It is noticeable that the mean steady-states for *in cis* and *in trans* models are qualitatively similar. In the case of standard deviations, we notice a substantial variance reduction in the model for the *in cis* system in comparison to the model of the *in trans* system. Intuitively, this is because we use one promoter instead of two and hence we have lower noise levels in comparison to two promoters. Mathematically, this reduction comes from the expression of the fmRNA which splits with a high probability into mGFP and sRNA relating the transcription probabilities of mGFP and sRNA, as well as the feedback of sRNA back on fmRNA.

The numerical simulations suggest that the *in cis* system is behaving similarly to the *in trans* systems in the context of the mean steady-state behaviour. Both systems allow tuning the range of the mean response to the inducer. However, the standard deviation at steady-state is much higher in the *in trans* systems in comparison to the *in cis* system.

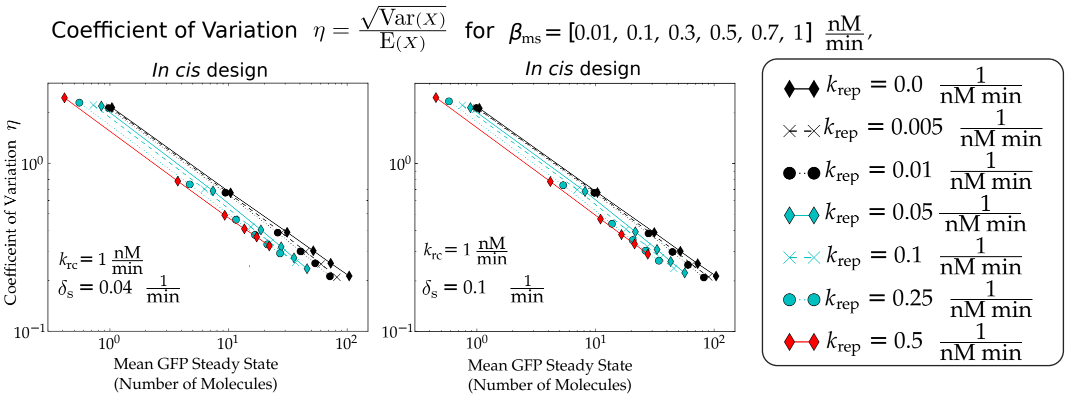


Figure S3: Comparison of the noise levels in the *in cis* for different parameter values. Data points for each line correspond to different values of $\beta_{\mathrm{ms}}$ from [0.01, 0.1, 0.3, 0.5, 0.7, 1] [nM/min]. Every line corresponds to a different repression sRNA-mRNA binding strength $k_{\mathrm{rep}}$. In the figure, $k_{\mathrm{rc}}$ stands for the ribozyme cleavage rate and $\delta_{\mathrm{s}}$ is the sRNA degradation rate.

##### S1.4.3 Reference tracking

In this subsection we study the tracking properties of our design. We perform simulations for the *in cis* system while varying $\beta_{\mathrm{ms}}$. We consider $\beta_{\mathrm{ms}} = \beta^0_{\mathrm{ms}}(1 + u(t))$, and set $u(t)$ as follows:

$$
\begin{aligned}
u(t) &= 1, && \text{if } t \in [0, T/8] \\
u(t) &= 2, && \text{if } t \in [T/4, 3T/8] \\
u(t) &= 3, && \text{if } t \in [T/2, 5T/8] \\
u(t) &= 10^{-2}(t - 3T/4), && \text{if } t \in [3T/4, 7T/8] \\
u(t) &= 0, && \text{otherwise}
\end{aligned}
$$

where $T = 3000$ [min] $= 50$ [h]. We perform the simulation in two settings: We set $k_{\mathrm{rc}} = 5$ [1/min] and simulate the model (S4), and simulate the model (S2) with $\beta_m = \beta_{\mathrm{ms}}$, $\beta_s = \beta_{\mathrm{ms}}$, effectively assuming that the ribozyme cleavage happens instantaneously and $k_{\mathrm{rc}} = \infty$, $\frac{k_{\mathrm{rc}}}{\delta_{\mathrm{m}}+k_{\mathrm{rc}}} = 1$. We perform simulations for different values of $\beta^0_{\mathrm{ms}}$, which we vary from $10^{-2}$ to 10 [1/min]. The GFP steady-state can be computed analytically, but we consider a series of approximations and obtain:

$$
[\mathrm{GFP}]^{\mathrm{ss}} \approx \frac{k_{\mathrm{t}}}{\delta_{\mathrm{p}}} \sqrt{\frac{\beta_{\mathrm{ms}}\delta_{\mathrm{s}}}{k_{\mathrm{rep}}\delta_{\mathrm{m}}}},
$$

which is valid if $\frac{k_{\mathrm{rc}}}{\delta_{\mathrm{m}}+k_{\mathrm{rc}}} \approx 1$, $4\frac{k_{\mathrm{rep}}\beta_{\mathrm{ms}}}{\delta_{\mathrm{m}}\delta_{\mathrm{s}}} \gg 1$. Therefore, the value $\frac{[\mathrm{GFP}]^{\mathrm{ss}}}{\sqrt{\beta_{\mathrm{ms}}}}$ remains constant at steady-state for constant parameters $k_{\mathrm{t}}$, $\delta_{\mathrm{p}}$, $\delta_{\mathrm{m}}$, $\delta_{\mathrm{s}}$, $k_{\mathrm{rep}}$ and changes in the "input" $\beta_{\mathrm{ms}}$, which can be used to compute "the ideal output" $y(t)$ given the input $u(t)$ of the model as follows $y(t) = \sqrt{1 + u(t)}$.

The trajectories with $k_{\mathrm{rc}} = \infty$ follow the ideal response very well, even in the case of the ramp. While there is a mismatch between the response and the ideal behaviour, this is barely visible and becomes indistinguishable for large values of $\beta_{\mathrm{ms}}$, which is consistent with the modelling. The trajectories with $k_{\mathrm{rc}} = 5$ have a significant mismatch with the ideal response, which indicates that engineering faster self-cleaving ribozymes can produce better filters.

We note that our results are similar in spirit to the antithetic control tracking [Briat et al., 2016, Agrawal et al., 2018], however, there are substantial differences with the major one being that we track a nonlinear function of the input.

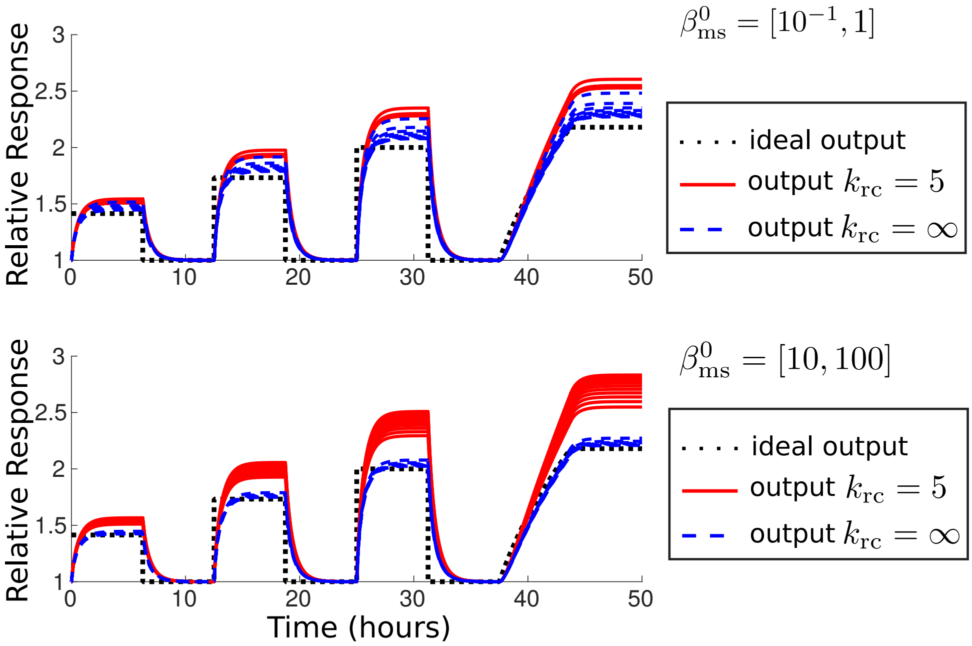


Figure S4: Trajectory tracking in the *in cis* model. We depict the relative GFP expression rate with respect to unperturbed expression rates ($u(t) = 1$ for all $t$). Assuming that the ribozyme cleavage happens instantaneously ($k_{\rm rc} = \infty$) allows for a near-perfect tracking of the desired output (computed as $\sqrt{1 + u(t)}$). In both simulations, there is a mismatch between the ideal output and the trajectory with $k_{\rm rc} = \infty$.

#### S1.5 Detailed theoretical analysis of the models

##### S1.5.1 Non-dimensionalisation

In order to simplify the analysis it is common to reduce the number of parameters in the system by introducing non-dimensional variables. We introduce the following non-dimensional variables and constants:

$$
\begin{aligned}
x_1 &= [\mathrm{mGFP^{tot}}] \left(\frac{\beta_{\rm ms}}{\delta_{\rm m}}\right)^{-1} & k_1 &= k_{\rm rep}\frac{\beta_{\rm ms}}{\delta_{\rm m}}, \\
x_2 &= [\mathrm{GFP}] \left(\frac{k_{\rm t}\beta_{\rm ms}}{\delta_{\rm m}\delta_{\rm p}}\right)^{-1} & k_2 &= \frac{k_{\rm rc}\delta_{\rm m}}{\delta_{\rm m} + k_{\rm rc}}, \\
x_3 &= [\mathrm{sRNA}] \left(\frac{\beta_{\rm ms}}{\delta_{\rm m}}\right)^{-1} & \delta_1 &= \frac{\delta_{\rm m} + k_{\rm rc}}{k_{\rm rc}}, \\
x_4 &= [\mathrm{fmRNA}] \left(\frac{\beta_{\rm ms}}{\delta_{\rm m} + k_{\rm rc}}\right)^{-1} & \delta_2 &= \frac{\delta_{\rm s} + k_{\rm rc}}{k_{\rm rc}}, \\
x_5 &= [\mathrm{sRNA:fmRNA}] \left(\frac{\beta_{\rm ms}}{\delta_{\rm m} + k_{\rm rc}}\right)^{-1} & \varepsilon &= \frac{1}{k_{\rm rc}}.
\end{aligned}
$$

This results in the following ordinary differential equation model:

$$
\begin{aligned}
\dot{x}_1 &= \delta_{\rm m}(1 - x_1) - k_1 x_3 x_1, \\
\dot{x}_2 &= x_1 - \delta_{\rm p} x_2, \\
\dot{x}_3 &= k_2(x_4 + x_5) - \delta_{\rm s} x_3 - k_1 x_3 x_1, \\
\varepsilon\dot{x}_4 &= \delta_1(1 - x_4) - \varepsilon k_1 x_3 x_4, \\
\varepsilon\dot{x}_5 &= -\delta_2 x_5 + \varepsilon k_1 x_3 x_4.
\end{aligned}
\tag{S7}
$$

This model is useful for two main reasons: The time-scales of the species are apparent, and the number of parameters is substantially reduced. We will interchangeably use the models (S7) and (S4) in our subsequent analysis.

##### S1.5.2 Trajectories of the system (S7) (and thus the system (S4)) are bounded

Specifically we have:

$$x_1 \in [0,1],\ x_2 \in \left[0, \frac{1}{\delta_{\mathrm{p}}}\right],\ x_3 \in \left[0, \frac{\delta_1 k_2}{\delta_{\mathrm{s}} \min(\delta_1,\delta_2)}\right],\ x_4 \in [0,1]\ x_5 \in \left[0, \frac{\delta_1}{\min(\delta_1,\delta_2)}\right].$$

For one dimensional models $\dot{z} = f(z)$ with $z \in \mathbb{R}$, we can derive simple bounds as follows. Assume there are functions $g(\cdot)$ and $h(\cdot)$ such that $g(z) \le f(z) \le h(z)$ for all $z \in \mathcal{D}$, where $\mathcal{D}$ is a compact set. Consider the trajectories of the systems $\dot{z} = g(z)$, $\dot{z} = f(z)$, $\dot{z} = h(z)$ originating from $z_0$, and denote them as $\phi_g(t, z_0)$, $\phi_f(t, z_0)$ $\phi_h(t, z_0)$, respectively. It is well-known that in this case we have $\phi_g(t, z_0) \le \phi_f(t, z_0) \le \phi_h(t, z_0)$ for all $t > 0$. Equipped with this simple result we can derive the bounds for the system (S7).

Consider first the lower bounds. By removing all nonnegative terms for nonnegative $x$ in the system (S7), we obtain the equations $\dot{x}_i = -x_i g_i(x)$, where $g_i(x)$ is nonnegative for all $x$. Therefore for every species, the trajectories of systems $\dot{x}_i = -x_i g_i(x)$ converge to zero for any nonnegative $x(t)$. Thus the lower bounds for all the species is zero.

The upper bounds for the states $x_1$ and $x_4$ are straightforward, since for nonnegative $x$ we can discard the nonlinear terms and thus obtain the vector fields $h_i$. Assuming that $x_1$ is at its maximum level, that is $x_1 = 1$, we obtain the upper bound for $x_2$. Now consider the equation for $x_4 + x_5$:

$$\frac{d}{dt}(x_4 + x_5) = \delta_1(1 - x_4) - \delta_2 x_5 \le \delta_1 - \min(\delta_1,\delta_2)(x_4 + x_5),$$

hence $x_4 + x_5 \le \frac{\min(\delta_1,\delta_2)}{\delta_1}$ and since $x_4$ is nonnegative, we obtain the upper bound for $x_5$. Now the bound for $x_3$ is also straightforward.

##### S1.5.3 Time-scale separation

With the growth of the ribozyme cleavage rate $k_{\mathrm{rc}}$, $\varepsilon$ is converging to 0. Furthermore, $k_{\mathrm{rc}}$ (which is the time-scale of the species $x_4$, $x_5$) is substantially larger than the time-scales of the species $x_1$, $x_2$, and $x_3$. Therefore, we can perform time-scale separation analysis by solving the last two equations in (S7) with $\varepsilon = 0$, which gives us $x_5 = 0$, $x_4 = 1$. Replacing the original variables gives us the reduced order model:

$$\begin{aligned}
\frac{d}{dt}[\mathrm{mGFP^{tot}}] &= \beta_{\mathrm{ms}} - \delta_{\mathrm{m}}[\mathrm{mGFP^{tot}}] - k_{\mathrm{rep}}[\mathrm{sRNA}][\mathrm{mGFP^{tot}}],\\
\frac{d}{dt}[\mathrm{GFP}] &= k_{\mathrm{t}}[\mathrm{mGFP^{tot}}] - \delta_{\mathrm{p}}[\mathrm{GFP}],\\
\frac{d}{dt}[\mathrm{sRNA}] &= \frac{k_{\mathrm{rc}}}{\delta_{\mathrm{m}} + k_{\mathrm{rc}}}\beta_{\mathrm{ms}} - \delta_{\mathrm{s}}[\mathrm{sRNA}] - k_{\mathrm{rep}}[\mathrm{sRNA}][\mathrm{mGFP^{tot}}].
\end{aligned} \tag{S8}$$

Therefore with $\varepsilon \to 0$, the trajectories of the model (S4) converge to the trajectories of (S8).

##### S1.5.4 Stability analysis

First, let us compute the steady-states. Assuming that $\frac{d}{dt}[\mathrm{mGFP^{tot}}] = 0$, we have:

$$[\mathrm{mGFP^{tot}}]^{ss} = \frac{\beta_{\mathrm{ms}}}{\delta_{\mathrm{m}} + k_{\mathrm{rep}}[\mathrm{sRNA}]^{ss}}.$$

Let $\alpha = \frac{k_{\mathrm{rc}}}{k_{\mathrm{rc}} + \delta_{\mathrm{m}}}$, then for sRNA steady-state we need to solve the following equation

$$0 = \alpha\beta_{\mathrm{ms}} - \delta_{\mathrm{s}}[\mathrm{sRNA}]^{ss} - \beta_{\mathrm{ms}}\frac{k_{\mathrm{rep}}[\mathrm{sRNA}]^{ss}}{\delta_{\mathrm{m}} + k_{\mathrm{rep}}[\mathrm{sRNA}]^{ss}},$$

which results in the quadratic equation

$$0 = \alpha\beta_{\mathrm{ms}}\delta_{\mathrm{m}} - k_{\mathrm{rep}}\delta_{\mathrm{s}}([\mathrm{sRNA}]^{ss})^2 - (\delta_{\mathrm{s}}\delta_{\mathrm{m}} + (1-\alpha)\beta_{\mathrm{ms}}k_{\mathrm{rep}})[\mathrm{sRNA}]^{ss}$$

with a unique positive root:

$$[\mathrm{sRNA}]^{ss} = -\frac{(\delta_{\mathrm{s}}\delta_{\mathrm{m}} + (1-\alpha)\beta_{\mathrm{ms}}k_{\mathrm{rep}})}{2k_{\mathrm{rep}}\delta_{\mathrm{s}}} + \\ + \frac{\sqrt{(\delta_{\mathrm{s}}\delta_{\mathrm{m}} + (1-\alpha)\beta_{\mathrm{ms}}k_{\mathrm{rep}})^2 + 4k_{\mathrm{rep}}\alpha\beta_{\mathrm{ms}}\delta_{\mathrm{m}}\delta_{\mathrm{s}}}}{2k_{\mathrm{rep}}\delta_{\mathrm{s}}}.$$

The steady-state for the GFP concentrations can be then computed as

$$[\mathrm{GFP}]^{\mathrm{ss}} = \frac{k_{\mathrm{t}}}{\delta_{\mathrm{p}}}\frac{\beta_{\mathrm{ms}}}{\delta_{\mathrm{m}} + k_{\mathrm{rep}}[\mathrm{sRNA}]^{ss}}.$$

The Jacobian of the vector field at the steady-state is computed as follows:

$$J(x) = \begin{pmatrix} -\delta_{\mathrm{m}} - k_{\mathrm{rep}}[\mathrm{sRNA}]^{\mathrm{ss}} & 0 & -k_{\mathrm{rep}}[\mathrm{mGFP}^{\mathrm{tot}}]^{\mathrm{ss}} \\ k_{\mathrm{t}} & -\delta_{\mathrm{p}} & 0 \\ -k_{\mathrm{rep}}[\mathrm{sRNA}]^{\mathrm{ss}} & 0 & -\delta_{\mathrm{s}} - k_{\mathrm{rep}}[\mathrm{mGFP}^{\mathrm{tot}}]^{\mathrm{ss}} \end{pmatrix},$$

The matrix $J(x)$ can be verified to be *strictly scaled diagonally dominant* (see for example [Varga, 1976] for definitions) and has negative diagonal terms for *all positive parameter values and nonnegative steady-states*. This implies that $J(x)$ is Hurwitz (all eigenvalues have negative real parts) and hence the system is locally exponentially stable (see for example [Sootla and Anderson, 2016, Sootla et al., 2017]), which means that all the trajectories from physically meaningful initial states asymptotically converge to the same steady-state.

Inspecting the sign-pattern of the Jacobian matrix $J(x)$ we notice that all feedback loops are positive. This means that the model (S8) is *monotone* [Hirsch et al., 2005] and hence has strong stability properties. In particular, if a monotone model has a unique locally asymptotically stable steady-state and the trajectories originating from a compact set $\mathcal{D}$ stay in $\mathcal{D}$, then this steady-state is attractive on $\mathcal{D}$ as well. Using these arguments we can show the following theoretical result:

**Theorem 1** *There exists a small enough value $\varepsilon$ such that for all $k_{\mathrm{rc}} > 1/\varepsilon$ and all positive parameter values, the system* (S4) *evolves on the compact set and all the trajectories are attracted to a unique attractive and locally stable steady-state.*

We will not prove this result formally, however, the proof is fairly straightforward using standard monotone systems tools and the results above.

##### S1.5.5 Frequency response analysis

In control theory, frequency domain analysis often reveals additional properties of the system. In order to perform the analysis we need to linearise the models around the steady-state. Consider the *in trans* model

$$\begin{aligned} \frac{d}{dt}[\mathrm{mGFP}] &= u - \delta_{\mathrm{m}}[\mathrm{mGFP}] - k_{\mathrm{rep}}[\mathrm{sRNA}][\mathrm{mGFP}], \\ \frac{d}{dt}[\mathrm{sRNA}] &= \beta_{\mathrm{s}} - \delta_{\mathrm{s}}[\mathrm{sRNA}] - k_{\mathrm{rep}}[\mathrm{sRNA}][\mathrm{mGFP}], \\ \frac{d}{dt}[\mathrm{GFP}] &= k_{\mathrm{t}}[\mathrm{mGFP}] - \delta_{\mathrm{p}}[\mathrm{GFP}], \\ y &= [\mathrm{GFP}], \end{aligned} \tag{S9}$$

where we treat the transcription rate of mGFP as an input to the model and GFP concentrations as the output of the model. In the *in cis* model we also treat GFP concentrations as the output

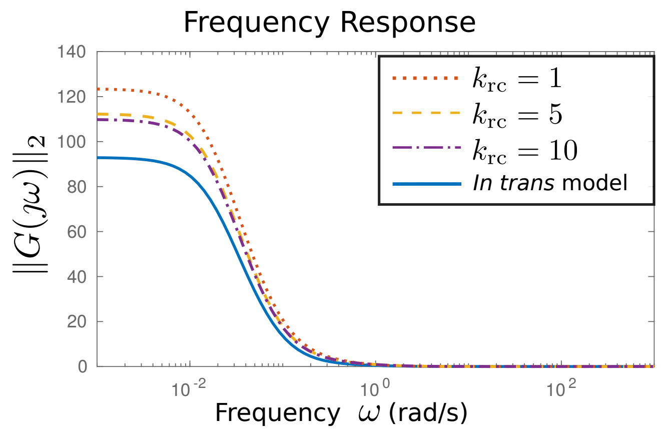


Figure S5: Frequency response of the linearised *in cis* design model for different values of $k_{\mathrm{rc}}$ and the linearised *in trans* design model.

of the model, but we treat the promoter activity as an input, which result in the following model:

$$\begin{aligned}\frac{d}{dt}[\mathrm{mGFP}] &= k_{\mathrm{rc}}[\mathrm{fmRNA}] - \delta_{\mathrm{m}}[\mathrm{mGFP}] - k_{\mathrm{rep}}[\mathrm{sRNA}][\mathrm{mGFP}] \\ \frac{d}{dt}[\mathrm{GFP}] &= k_{\mathrm{t}}([\mathrm{mGFP}] + [\mathrm{fmRNA}]) - \delta_{\mathrm{p}}[\mathrm{GFP}], \\ \frac{d}{dt}[\mathrm{sRNA}] &= k_{\mathrm{rc}}([\mathrm{sRNA}:\mathrm{fmRNA}] + [\mathrm{fmRNA}]) - \\ &\quad - \delta_{\mathrm{s}}[\mathrm{sRNA}] - k_{\mathrm{rep}}[\mathrm{sRNA}]([\mathrm{mGFP}] + [\mathrm{fmRNA}]) \\ \frac{d}{dt}[\mathrm{fmRNA}] &= u - \delta_{\mathrm{m}}[\mathrm{fmRNA}] - k_{\mathrm{rc}}[\mathrm{fmRNA}] - \\ &\quad - k_{\mathrm{rep}}[\mathrm{sRNA}][\mathrm{fmRNA}] \\ \frac{d}{dt}[\mathrm{sRNA}:\mathrm{fmRNA}] &= k_{\mathrm{rep}}[\mathrm{sRNA}][\mathrm{fmRNA}] - \\ &\quad - \delta_{\mathrm{s}}[\mathrm{sRNA}:\mathrm{fmRNA}] - k_{\mathrm{rc}}[\mathrm{sRNA}:\mathrm{fmRNA}], \\ y &= [\mathrm{GFP}].\end{aligned} \tag{S10}$$

Note that in the *in cis* model, the input affects both mGFP and fmRNA. This interpretation allows us to treat the model in the input-output framework, which allows us to perform the frequency domain analysis of the model. In order to do so we need to linearise the model about the steady-state. Take a nonlinear model:

$$\begin{aligned} z &= f(z,u) \\ y &= h(z) \end{aligned}$$

with the steady-state $z^*$ taken for a particular value of $u$ denoted as $u^*$. We will study the deviations around the steady-state:

$$\begin{aligned} \Delta z &= A\Delta z + B\Delta u, \\ \Delta y &= C\Delta z, \end{aligned}$$

where the matrices $A = \partial f(z,u)/\partial z(z^*,u^*)$, $B = \partial f(z,u)/\partial u(z^*,u^*)$ and $C = \partial h(z)/\partial z(z^*)$ are the first terms of the Taylor expansion of $f(z,u)$ and $h(z)$ about $z^*$, $u^*$. Taking the Laplace transform of the linear model results in the relation

$$\Delta Y(s) = \underbrace{C(sI - A)^{-1}B}_{G(s)} \Delta U(s)$$

and enables the frequency domain analysis. In particular, the value $\|G(\jmath\omega)\|_2$, where $\jmath$ is the complex identity and $\omega$ is a real number, signifies how the system reacts to a sinusoidal input with the frequency $\omega$ ($u(t) = \sin(\omega t)$). If $\|G(\jmath\omega)\|_2 > 1$ then the signal $u(t)$ is amplified and if $\|G(\jmath\omega)\| < 1$ then the signal $u(t)$ is attenuated. We plot the function $\|G(\jmath\omega)\|_2$ in Figure S5, which indicates that both designs act as *low-pass filters*, which means that the systems react to constant inputs (and inputs with low frequency), but attenuate high-frequency signals such as noise, which both are desirable properties.

### S2 Design and modelling of the enhanced TetR autorepressor

#### S2.1 The designs

As the one of the applications of our module we envision tuning the strength of a feedback interconnection. We take the classical TetR autorepressor design. We compare two ways of tuning the feedback strength in the TetR autorepressor design: with the sRNA production outside the loop (*in trans* design) and with the sRNA production inside the loop (*in cis* design), see Figure 6 in the main text. Since the modelling process is similar to the filter case, we only sketch the main theoretical results and model derivations.

##### S2.1.1 Model of the *in trans* design

Our modelling procedure is fairly similar to the filter case, with a few exceptions: We employ a $P_{\mathrm{tet}}$ promoter and place downstream of $P_{\mathrm{tet}}$ a fused *gfp* with *tetR*, which represses the activity of the $P_{\mathrm{tet}}$ promoter. In *in trans* design, we consider the following chemical reactions:

$$\boxed{\begin{array}{lll} \emptyset \xrightarrow{\beta_{\mathrm{t}}} \mathrm{mTetR} & \emptyset \xrightarrow{\beta_{\mathrm{s}}} \mathrm{sRNA} & \\ \mathrm{mTetR} \xrightarrow{\mathrm{k_t}} \mathrm{mTetR} + \mathrm{TetR} & \mathrm{sRNA} + \mathrm{mTetR} \xrightarrow{\mathrm{k_{rep}}} \emptyset & \\ \mathrm{mTetR} \xrightarrow{\delta_{\mathrm{m}}} \emptyset & \mathrm{sRNA} \xrightarrow{\delta_{\mathrm{s}}} \emptyset & \mathrm{TetR} \xrightarrow{\delta_{\mathrm{t}}} \emptyset. \end{array}} \tag{S11}$$

where $\beta_{\mathrm{t}} = \beta_{\mathrm{s}} = \beta_{\mathrm{tx}}$ and the promoter activity is modelled as follows:

$$\beta_{\mathrm{tx}} = \frac{K_0}{1 + K_1 + 2K_2\dfrac{[\mathrm{TetR}]}{1 + [\mathrm{aTc}]/K_D} + \left(K_2\dfrac{[\mathrm{TetR}]}{1 + [\mathrm{aTc}]/K_D}\right)^2}.$$

Note that we assume that both sRNA, and TetR are under the identical $P_{\mathrm{tet}}$ promoter. We can naturally place sRNA under a different inducible promoter (for example, arabinose inducible $P_{\mathrm{bad}}$ promoter), however, in this case sRNA is produced in the open-loop. The closed-loop production was guided by the subsequent comparison to the *in cis* design.

We do not model GFP production since GFP is fused with TetR and serves as a reporter on TetR. We model the $P_{\mathrm{tet}}$ promoter as described above and assume that the rest of the interactions follow mass action kinetics. We obtain the following ODE model:

$$\boxed{\begin{aligned} \frac{d}{dt}[\mathrm{mTetR}] &= \beta_{\mathrm{t}} - \delta_{\mathrm{m}}[\mathrm{mTetR}] - \mathrm{k_{rep}}[\mathrm{sRNA}][\mathrm{mTetR}] \\ \frac{d}{dt}[\mathrm{sRNA}] &= \beta_{\mathrm{s}} - \delta_{\mathrm{s}}[\mathrm{sRNA}] - k_{\mathrm{rep}}[\mathrm{sRNA}][\mathrm{mTetR}] \\ \frac{d}{dt}[\mathrm{TetR}] &= k_{\mathrm{t}}([\mathrm{mTetR}]) - \delta_{\mathrm{t}}[\mathrm{TetR}], \end{aligned}} \tag{S12}$$

##### S2.1.2 Model of the *in cis* design

We model the *in cis* design similarly and consider the following chemical reactions

$$
\begin{array}{|ll|}
\hline
\emptyset \xrightarrow{\beta_{\mathrm{ms}}} \mathrm{fmRNA} & \\
\mathrm{fmRNA} \xrightarrow{k_{\mathrm{rc}}} \mathrm{mTetR} + \mathrm{sRNA} & \mathrm{sRNA : fmRNA} \xrightarrow{k_{\mathrm{rc}}} \mathrm{sRNA} \\
\mathrm{sRNA} + \mathrm{fmRNA} \xrightarrow{k_{\mathrm{rep}}} \mathrm{sRNA : fmRNA} & \mathrm{sRNA} + \mathrm{mTetR} \xrightarrow{\mathrm{k_{rep}}} \emptyset \\
\mathrm{mTetR} \xrightarrow{\mathrm{k_t}} \mathrm{mTetR} + \mathrm{TetR} & \mathrm{fmRNA} \xrightarrow{k_{\mathrm{t}}} \mathrm{fmRNA} + \mathrm{TetR} \\
\mathrm{mTetR} \xrightarrow{\delta_{\mathrm{m}}} \emptyset & \mathrm{sRNA} \xrightarrow{\delta_{\mathrm{s}}} \emptyset, \ \mathrm{TetR} \xrightarrow{\delta_{\mathrm{t}}} \emptyset. \\
\hline
\end{array}
\tag{S13}
$$

where $\beta_{\mathrm{ms}} = \beta_{\mathrm{tx}}$ and the promoter activity is modelled as follows:

$$\beta_{\mathrm{tx}} = \frac{K_0}{1 + K_1 + 2K_2 \frac{[\mathrm{TetR}]}{1 + [\mathrm{aTc}]/K_D} + \left(K_2 \frac{[\mathrm{TetR}]}{1 + [\mathrm{aTc}]/K_D}\right)^2}.$$

We obtain the following ODE model:

$$
\boxed{
\begin{aligned}
\frac{d}{dt}[\mathrm{mTetR}] &= k_{\mathrm{rc}}[\mathrm{fmRNA}] - \delta_{\mathrm{m}}[\mathrm{mTetR}] - \mathrm{k_{rep}}[\mathrm{sRNA}][\mathrm{mTetR}] \\
\frac{d}{dt}[\mathrm{fmRNA}] &= \beta_{\mathrm{ms}} - \delta_{\mathrm{m}}[\mathrm{fmRNA}] - k_{\mathrm{rc}}[\mathrm{fmRNA}] - \\
&\quad - k_{\mathrm{rep}}[\mathrm{sRNA}][\mathrm{fmRNA}] \\
\frac{d}{dt}[\mathrm{sRNA : fmRNA}] &= k_{\mathrm{rep}}[\mathrm{sRNA}][\mathrm{fmRNA}] - \delta_{\mathrm{s}}[\mathrm{sRNA : fmRNA}] - \\
&\quad - k_{\mathrm{rc}}[\mathrm{sRNA : fmRNA}] \\
\frac{d}{dt}[\mathrm{sRNA}] &= k_{\mathrm{rc}}([\mathrm{sRNA : fmRNA}] + [\mathrm{fmRNA}]) - \delta_{\mathrm{s}}[\mathrm{sRNA}] - \\
&\quad - k_{\mathrm{rep}}[\mathrm{sRNA}]([\mathrm{mTetR}] + [\mathrm{fmRNA}]) \\
\frac{d}{dt}[\mathrm{TetR}] &= k_{\mathrm{t}}([\mathrm{mTetR}] + [\mathrm{fmRNA}]) - \delta_{\mathrm{t}}[\mathrm{TetR}],
\end{aligned}
}
\tag{S14}
$$

As in the case of the filter, mTetR and fmRNA are under the same promoter. We can also apply the time-scale separation arguments provided that the ribozyme cleavage rate $k_{\mathrm{rc}}$ is large enough and reach the following reduced order model:

$$
\boxed{
\begin{aligned}
\frac{d}{dt}[\mathrm{mTetR}] &= \beta_{\mathrm{ms}} - \delta_{\mathrm{m}}[\mathrm{mTetR}] - \mathrm{k_{rep}}[\mathrm{sRNA}][\mathrm{mTetR}] \\
\frac{d}{dt}[\mathrm{sRNA}] &= \frac{k_{\mathrm{rc}}}{\delta_{\mathrm{m}} + k_{\mathrm{rc}}}\beta_{\mathrm{ms}} - \delta_{\mathrm{s}}[\mathrm{sRNA}] - k_{\mathrm{rep}}[\mathrm{sRNA}][\mathrm{mTetR}] \\
\frac{d}{dt}[\mathrm{TetR}] &= k_{\mathrm{t}}[\mathrm{mTetR}] - \delta_{\mathrm{t}}[\mathrm{TetR}].
\end{aligned}
}
\tag{S15}
$$

Analytic computation of the steady-states in this case is more involved and we were not able to obtain an analytic formula for either of the designs. However, we can predict the changes in the steady-state by using the steady-state computations for the filter case. For example, if the ribozyme cleavage rate is large enough then we need to find the solution to the following two equations:

$$
\begin{aligned}
\beta_{\mathrm{tx}} = f_1([\mathrm{TetR}]) &= \frac{\mathrm{K_0}}{1 + \mathrm{K_1} + 2\mathrm{K_2}\frac{[\mathrm{TetR}]}{1 + [\mathrm{aTc}]/\mathrm{K_D}} + \left(\mathrm{K_2}\frac{[\mathrm{TetR}]}{1 + [\mathrm{aTc}]/\mathrm{K_D}}\right)^2}, \\
[\mathrm{TetR}] = f_2(\beta_{\mathrm{tx}}) &= \frac{k_{\mathrm{t}}}{\delta_{\mathrm{p}}} \frac{2\beta_{\mathrm{tx}}/\delta_{\mathrm{m}}}{1 + \sqrt{1 + 4\frac{k_{\mathrm{rep}}\beta_{\mathrm{tx}}}{\delta_{\mathrm{m}}\delta_{\mathrm{s}}}}}.
\end{aligned}
$$

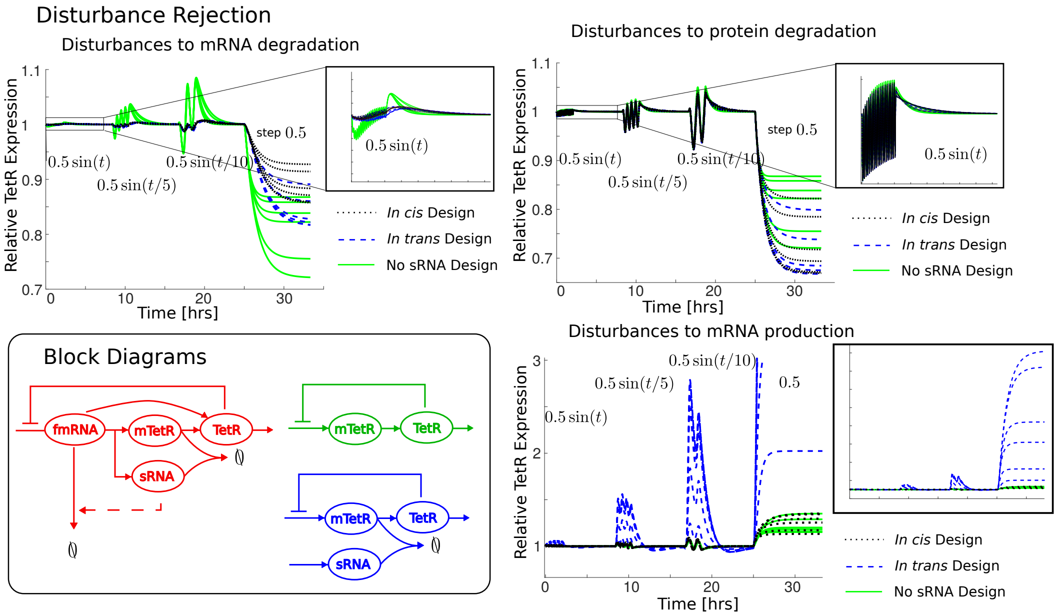


Figure S6: Disturbances to different parameters in the models. The double feedback loop in the *in cis* design allows us to effectively reject various disturbances. The *in trans* design amplifies the disturbances in the mRNA production since sRNA is produced under the different promoter. The classical autorepressor (without sRNA tuning) amplifies the disturbances in the mRNA degradation as these dynamics are not directly controlled by the transcription factor TetR.

The curve $\beta_{\text{tx}} = f_1([\text{TetR}])$ is monotonically decreasing in [TetR], while the curve $[\text{TetR}] = \text{f}_2(\beta_{\text{tx}})$ is monotonically increasing in $\beta_{\text{tx}}$ for all nonnegative parameter values, therefore there is a unique steady-state. Furthermore, the curve $f_2$ is decreasing with respect to $k_{\text{rep}}$, therefore the intersection point between $f_1$ and $f_2$ will increase in [TetR] and decrease in $\beta_{\text{tx}}$. We can deduce similar dependencies with respect to other parameters. This means that there is a unique steady-state in this system and bistability can be ruled out.

#### S2.2 Theoretical comparison of the designs

##### S2.2.1 Parameter values

We take the parameter values fitted for the *in cis* filter studied above. We set $K_0 = 2.26$ [nM/min], $K_1 = 1.054$ [dimensionless], $K_2 = 18.46$ [dimensionless], $K_D = 0.1182$ [nM], $\delta_{\text{s}} = 0.0482$ [1/min], $\delta_{\text{m}} = 0.2476$ [1/min], $\delta_{\text{p}} = 0.0234$ [1/min], $k_{\text{rc}} = 1$ [1/min], $k_{\text{t}} = 1$ [1/min]. We will sweep through the repression rate $k_{\text{rep}}$ and aTc concentrations. In particular, we use the values $[0, 0.005, 0.01, 0.05, 0.1, 0.25, 0.5]$ [1/ (nM min)] for $k_{\text{rep}}$ and the values $[0.1, 1, 5, 10, 50, 100]$ [nM] for aTc concentrations.

##### S2.2.2 Disturbance rejection

We perform disturbance rejection analysis for the deterministic model. First, we set up the disturbance signals as follows:

$$\begin{aligned} w(t) &= 0.5\sin(t), && \text{if } t \in [0, T/16],\\ w(t) &= 0.5\sin(t/5), && \text{if } t \in [T/4, 5T/16],\\ w(t) &= 0.5\sin(t/10), && \text{if } t \in [T/2, 9T/16],\\ w(t) &= 0.5, && \text{if } t \in [3T/4, T],\\ w(t) &= 0, && \text{otherwise,} \end{aligned}$$

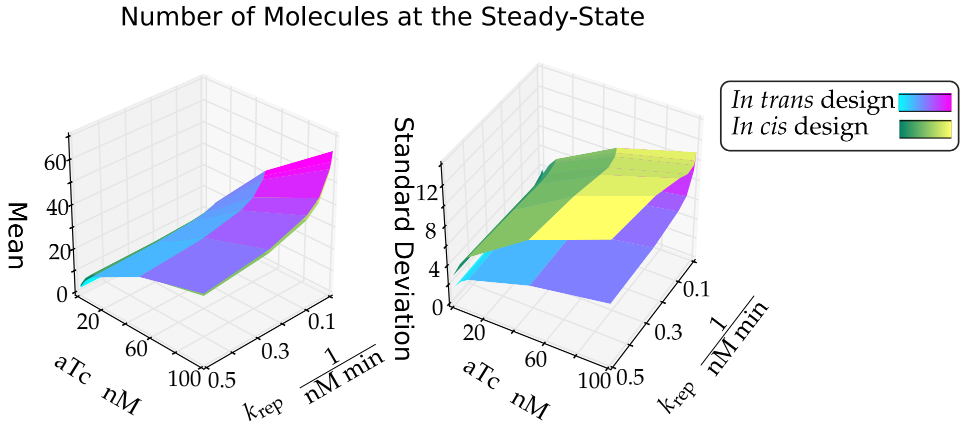


Figure S7: Model Comparison of the *in cis* and *in trans* feedback strength tuning. Statistics of GFP molecule number at $t = 2000$ [min] while performing a sweep over aTc concentrations and $k_{\text{rep}}$. The mean number of molecules are similar for the two designs, while the standard deviations significantly differ.

where $T$ is the simulation time set to 2000 [min] or 33.33 [hrs]. We choose to disturb three parameters TetR maximal production rate $\beta_{\text{tx}}$, the degradation rate of the mRNA $\delta_{\text{m}}$ and the degradation rate of TetR $\delta_{\text{p}}$ as follows:

$$\begin{aligned}
\beta_{\text{tx}} &= \beta^0_{\text{tx}}(1 + w(t)),\\
\delta_{\text{m}} &= \delta^0_{\text{m}}(1 + w(t)),\\
\delta_{\text{p}} &= \delta^0_{\text{p}}(1 + w(t)).
\end{aligned}$$

We note that in the *in cis* design, the disturbance affecting mRNA degradation rate also affects the functional mRNA degradation rate. We plot the simulation results in Figure S6. It is noticeable that the *in cis* design is less affected by the disturbances than the *in trans* design, which is especially noticeable for disturbances in the mRNA production rate. This is because the sRNA is produced under the same promoter as the mRNA of TetR in the *in cis* design, but also because the *in cis* design contains a feedback component. Furthermore, the system in the *in cis* design responds better to disturbances in the mRNA production and the degradation than the classical autorepressor design. Response to the disturbances $w(t)$ in the protein degradation is largely similar for all designs, but as Figure S6 suggests the classical autorepressor represses the step disturbances in $\delta_{\text{p}}$ slightly better. Overall the disturbance rejection properties in the *in cis* design and the classical autorepressor design (no sRNA) are quite similar, except for the disturbances in the mRNA degradation rate, where the *in cis* design is favourable. Additionally, using the *in cis* design we can adjust the mean response by changing the sRNA repression rate. Furthermore, the results in the main text indicate that the noise characteristics of the *in cis* are more favourable than the classical autorepressor design. We provide additional simulation data in the following.

###### S2.2.3 Mean, standard deviation and coefficient of variation at the steady states

We vary the repression strength $k_{\text{rep}}$, and aTc levels in our simulations, results of which are depicted in Figure S7. For the outside of the loop tuning (*in trans* design), we adjust the nominal production level of sRNA to match with the production level of the inside the loop tuning (*in cis* design).

Similarly to the filter design, the mean steady-state levels do not vary significantly. The major difference between the designs is visible in the noise characteristics. The sRNA tuning mechanism in *in cis* design attenuates noise for a wider range of parameters in comparison to the mechanism in *in trans* design. As for the case of the filter, in the *in cis* design the bursts in the production of mRNA are repressed by the bursts in the production of sRNA, which happen simultaneously with the bursts in the mRNA production.

### S3 Supplementary tables

#### S3.1 Coefficient of variation data

In this subsection we present the numerical values for Figures 5 and Figure 6 in the main text. Some values were omitted from the figure to improve readability.

Table S2: Mean and coefficient of variation rounded to the second significant digit in the filter design experiments. The comparable mean GFP steady-states are colour-coded.

*In cis* design

| $k_{\text{rep}}$[1/(nM min)] | Statistics | Production rate $\beta$ [nM/min] | | | | | |
|---|---|---|---|---|---|---|---|
| | | 0.01 | 0.1 | 0.3 | 0.5 | 0.7 | 1 |
| 0 | $\mathbb{E}(X)$ | 1.04 | 10.40 | 31.22 | 51.86 | 72.74 | 104.14 |
| | $\eta$ | 2.15 | 0.67 | 0.39 | 0.30 | 0.25 | 0.21 |
| 0.005 | $\mathbb{E}(X)$ | 0.99 | 9.92 | 28.04 | 44.77 | 60.39 | 81.98 |
| | $\eta$ | 2.16 | 0.67 | 0.39 | 0.30 | 0.25 | 0.21 |
| 0.01 | $\mathbb{E}(X)$ | 0.98 | 9.42 | 26.00 | 40.50 | 53.78 | 71.60 |
| | $\eta$ | 2.13 | 0.67 | 0.39 | 0.30 | 0.25 | 0.21 |
| 0.05 | $\mathbb{E}(X)$ | 0.85 | 7.40 | 18.76 | 27.79 | 35.65 | 45.96 |
| | $\eta$ | 2.19 | 0.68 | 0.40 | 0.32 | 0.27 | 0.24 |
| 0.1 | $\mathbb{E}(X)$ | 0.72 | 6.27 | 15.38 | 22.46 | 28.61 | 36.74 |
| | $\eta$ | 2.22 | 0.70 | 0.42 | 0.34 | 0.29 | 0.26 |
| 0.25 | $\mathbb{E}(X)$ | 0.55 | 4.71 | 11.54 | 16.72 | 21.23 | 27.20 |
| | $\eta$ | 2.29 | 0.75 | 0.46 | 0.37 | 0.34 | 0.29 |
| 0.5 | $\mathbb{E}(X)$ | 0.41 | 3.73 | 9.29 | 13.61 | 17.28 | 22.07 |
| | $\eta$ | 2.45 | 0.78 | 0.49 | 0.41 | 0.36 | 0.32 |

*In trans* design

| $k_{\text{rep}}$[1/(nM min)] | Statistics | 0.01 | 0.1 | 0.3 | 0.5 | 0.7 | 1 |
|---|---|---|---|---|---|---|---|
| 0.0 | $\mathbb{E}(X)$ | 1.03 | 10.44 | 31.17 | 51.93 | 72.73 | 104.22 |
| | $\eta$ | 2.14 | 0.67 | 0.39 | 0.30 | 0.25 | 0.21 |
| 0.005 | $\mathbb{E}(X)$ | 1.03 | 9.99 | 28.13 | 44.22 | 58.95 | 79.17 |
| | $\eta$ | 2.13 | 0.67 | 0.40 | 0.31 | 0.27 | 0.23 |
| 0.01 | $\mathbb{E}(X)$ | 1.04 | 9.66 | 25.89 | 39.99 | 51.91 | 68.11 |
| | $\eta$ | 2.12 | 0.68 | 0.40 | 0.32 | 0.28 | 0.24 |
| 0.05 | $\mathbb{E}(X)$ | 1.00 | 8.20 | 19.03 | 27.21 | 33.84 | 42.37 |
| | $\eta$ | 2.15 | 0.72 | 0.47 | 0.40 | 0.36 | 0.37 |
| 0.1 | $\mathbb{E}(X)$ | 0.99 | 7.37 | 16.12 | 22.30 | 27.51 | 33.97 |
| | $\eta$ | 2.14 | 0.76 | 0.52 | 0.46 | 0.42 | 0.39 |
| 0.25 | $\mathbb{E}(X)$ | 0.94 | 6.37 | 13.09 | 17.64 | 21.45 | 26.18 |
| | $\eta$ | 2.19 | 0.84 | 0.62 | 0.55 | 0.52 | 0.49 |
| 0.5 | $\mathbb{E}(X)$ | 0.94 | 5.82 | 11.45 | 15.40 | 18.50 | 22.40 |
| | $\eta$ | 2.18 | 0.89 | 0.69 | 0.62 | 0.60 | 0.57 |

Coefficient of variation and means for the feedback strength tuning simulations. The values correspond to Figure 6 in the main text. Some values were omitted from the figure to improve readability.

Table S3: Mean and coefficient of variation data rounded to the second significant digit in the feedback strength tuning simulations. The comparable mean GFP steady-states are colour-coded.

*In cis* design

| $k_{\text{rep}}[1/(\text{nM min})]$ | Statistics | aTc levels [nM] 0.01 | 0.1 | 0.3 | 0.5 | 0.7 | 1 |
|---|---|---|---|---|---|---|---|
| 0.0 | $\mathbb{E}(X)$ | 3.78 | 5.13 | 10.73 | 15.82 | 41.67 | 63.71 |
| | $\eta$ | 1.00 | 0.66 | 0.42 | 0.34 | 0.21 | 0.17 |
| 0.005 | $\mathbb{E}(X)$ | 3.60 | 5.03 | 10.49 | 15.48 | 39.65 | 59.29 |
| | $\eta$ | 0.98 | 0.66 | 0.42 | 0.33 | 0.20 | 0.17 |
| 0.01 | $\mathbb{E}(X)$ | 3.53 | 4.93 | 10.32 | 15.08 | 38.17 | 56.40 |
| | $\eta$ | 0.97 | 0.65 | 0.41 | 0.33 | 0.20 | 0.16 |
| 0.05 | $\mathbb{E}(X)$ | 3.03 | 4.49 | 9.29 | 13.46 | 32.23 | 45.97 |
| | $\eta$ | 0.89 | 0.62 | 0.39 | 0.31 | 0.19 | 0.16 |
| 0.1 | $\mathbb{E}(X)$ | 2.74 | 4.16 | 8.58 | 12.38 | 29.07 | 40.95 |
| | $\eta$ | 0.85 | 0.60 | 0.38 | 0.31 | 0.20 | 0.17 |
| 0.25 | $\mathbb{E}(X)$ | 2.33 | 3.63 | 7.62 | 10.93 | 25.15 | 34.70 |
| | $\eta$ | 0.80 | 0.57 | 0.37 | 0.30 | 0.20 | 0.18 |
| 0.5 | $\mathbb{E}(X)$ | 2.07 | 3.27 | 6.90 | 9.94 | 22.58 | 30.73 |
| | $\eta$ | 0.77 | 0.55 | 0.37 | 0.30 | 0.22 | 0.20 |

*In trans* design

| $k_{\text{rep}}[1/(\text{nM min})]$ | Statistics | aTc levels [nM] 0.01 | 0.1 | 0.3 | 0.5 | 0.7 | 1 |
|---|---|---|---|---|---|---|---|
| 0.0 | $\mathbb{E}(X)$ | 3.74 | 5.12 | 10.74 | 15.82 | 41.66 | 63.77 |
| | $\eta$ | 1.00 | 0.67 | 0.42 | 0.34 | 0.21 | 0.17 |
| 0.005 | $\mathbb{E}(X)$ | 3.64 | 5.05 | 10.54 | 15.44 | 39.52 | 58.77 |
| | $\eta$ | 0.99 | 0.67 | 0.42 | 0.34 | 0.21 | 0.17 |
| 0.01 | $\mathbb{E}(X)$ | 3.53 | 4.98 | 10.38 | 15.17 | 37.87 | 55.37 |
| | $\eta$ | 0.97 | 0.66 | 0.42 | 0.34 | 0.21 | 0.18 |
| 0.05 | $\mathbb{E}(X)$ | 3.13 | 4.68 | 9.57 | 13.65 | 31.54 | 43.99 |
| | $\eta$ | 0.95 | 0.66 | 0.43 | 0.35 | 0.24 | 0.22 |
| 0.1 | $\mathbb{E}(X)$ | 2.94 | 4.50 | 9.08 | 12.81 | 28.49 | 38.84 |
| | $\eta$ | 0.96 | 0.67 | 0.45 | 0.37 | 0.28 | 0.26 |
| 0.25 | $\mathbb{E}(X)$ | 2.78 | 4.29 | 8.48 | 11.79 | 25.05 | 33.11 |
| | $\eta$ | 1.00 | 0.70 | 0.49 | 0.42 | 0.35 | 0.35 |
| 0.5 | $\mathbb{E}(X)$ | 2.74 | 4.21 | 8.17 | 11.30 | 23.09 | 29.76 |
| | $\eta$ | 1.05 | 0.75 | 0.53 | 0.47 | 0.41 | 0.42 |

#### S3.2 Plasmids

List of plasmids used in the study.

Table S4: List of plasmids used for this study. All plasmids are based on pBbS2a-RFP [Lee et al., 2011] with the indicated modifications.

| Plasmids | Description | Reference |
|---|---|---|
| pBbS2a-RFP | Low copy number plasmid (pSC101 origin) – rfp under $P_{\text{tet}}$ promoter – Ampicillin resistance. | Shared bt Prof J. Keasling [Lee et al., 2011] |
| pND113 | *sfgfp* under $P_{\text{tet}}$ promoter | This work |
| pND149 | *sfgfp* in place of *rfp* – sRNA (25 nucleotides long TBS) under $P_{\text{tet}}$ promoter | This work |
| pND179 | *sfgfp* in place of *rfp* – HHR9 - sRNA (25 nucleotides long TBS) under $P_{\text{tet}}$ promoter | This work |
| pND218 | *sfgfp* in place of *rfp* – HHR9 - sRNA (30 nucleotides long TBS) under $P_{\text{tet}}$ promoter | This work |
| pND219 | *sfgfp* in place of *rfp* – HHR9 - sRNA (22 nucleotides long TBS) under $P_{\text{tet}}$ promoter | This work |
| pND221 | *sfgfp* in place of *rfp* – HHR9 - sRNA (25 nucleotides long TBS) under $P_{\text{tet}}$ promoter | This work |

### References


[Agrawal et al., 2018] Agrawal, D., Tang, X., Westbrook, A., Marshall, R., Maxwell, C., Lucks, J., Noireaux, V., Beisel, C., Dunlop, M. and Franco, E. (2018). Mathematical modeling of RNA-based architectures for closed loop control of gene expression. ACS synthetic biology *7*, 1219–1228.

[Briat et al., 2016] Briat, C., Gupta, A. and Khammash, M. (2016). Antithetic integral feedback ensures robust perfect adaptation in noisy biomolecular networks. Cell systems *2*, 15–26.

[Chen et al., 2015] Chen, H., Shiroguchi, K., Ge, H. and Xie, X. S. (2015). Genome-wide study of mRNA degradation and transcript elongation in Escherichia coli. Molecular systems biology *11*, 781.

[Gillespie, 1977] Gillespie, D. (1977). Exact stochastic simulation of coupled chemical reactions. The journal of physical chemistry *81*, 2340–2361.

[Hammann et al., 2012] Hammann, C., Luptak, A., Perreault, J. and De La Peña, M. (2012). The ubiquitous hammerhead ribozyme. RNA *18*, 871–885.

[Hirsch et al., 2005] Hirsch, M., Smith, H. et al. (2005). Monotone dynamical systems. Handbook of differential equations: ordinary differential equations *2*, 239–357.

[Hussein and Lim, 2012] Hussein, R. and Lim, H. (2012). Direct comparison of small RNA and transcription factor signaling. Nucleic acids research *40*, 7269–7279.

[Kelly et al., 2018] Kelly, C., Harris, A., Steel, H., Hancock, E., Heap, J. and Papachristodoulou, A. (2018). Synthetic negative feedback circuits using engineered small RNAs. Nucleic acids research *46*, 9875–9889.

[Kennell and Riezman, 1977] Kennell, D. and Riezman, H. (1977). Transcription and translation initiation frequencies of the Escherichia coli lac operon. Journal of molecular biology *114*, 1–21.

[Lee et al., 2011] Lee, T., Krupa, R., Zhang, F., Hajimorad, M., Holtz, W., Prasad, N., Lee, S. and Keasling, J. (2011). BglBrick vectors and datasheets: a synthetic biology platform for gene expression. Journal of biological engineering *5*, 12.

[Liang et al., 1999] Liang, S.-T., Bipatnath, M., Xu, Y.-C., Chen, S.-L., Dennis, P., Ehrenberg, M. and Bremer, H. (1999). Activities of constitutive promoters in Escherichia coli1. Journal of molecular biology *292*, 19–37.

[McCullen et al., 2010] McCullen, C., Benhammou, J., Majdalani, N. and Gottesman, S. (2010). Mechanism of positive regulation by DsrA and RprA small noncoding RNAs: pairing increases translation and protects rpoS mRNA from degradation. Journal of bacteriology *192*, 5559–5571.

[Perreault et al., 2011] Perreault, J., Weinberg, Z., Roth, A., Popescu, O., Chartrand, P., Ferbeyre, G. and Breaker, R. (2011). Identification of hammerhead ribozymes in all domains of life reveals novel structural variations. PLoS computational biology *7*, e1002031.

[Sootla and Anderson, 2016] Sootla, A. and Anderson, J. (2016). On existence of solutions to structured Lyapunov inequalities. In Proc of Am Control Conf pp. 7013–7018, IEEE.

[Sootla et al., 2017] Sootla, A., Zheng, Y. and Papachristodoulou, A. (2017). Block-Diagonal Solutions to Lyapunov Inequalities and Generalisations of Diagonal Dominance. In Proc IEEE Conf Decision Control.

[Steel et al., 2017] Steel, H., Harris, A., Hancock, E., Kelly, C. and Papachristodoulou, A. (2017). Frequency domain analysis of small non-coding RNAs shows summing junction-like behaviour. In Proc Conf Decision Control pp. 5328–5333, IEEE.

[Tamsir et al., 2011] Tamsir, A., Tabor, J. and Voigt, C. (2011). Robust multicellular computing using genetically encoded NOR gates and chemical wires. Nature *469*, 212–215.

[Varga, 1976] Varga, R. (1976). On recurring theorems on diagonal dominance. Linear Algebra and its Applications *13*, 1–9.

[Weinberg et al., 2015] Weinberg, Z., Kim, P., Chen, T., Li, S., Harris, K., Lünse, C. and Breaker, R. (2015). New classes of self-cleaving ribozymes revealed by comparative genomics analysis. Nature chemical biology *11*, 606.

[Zhou et al., 2011] Zhou, Y., Liepe, J., Sheng, X., Stumpf, M. and Barnes, C. (2011). GPU accelerated biochemical network simulation. Bioinformatics *27*, 874–876.
